# ALOHA: Aggregated local extrema splines for high-throughput dose-response analysis

**DOI:** 10.1101/2021.03.29.437588

**Authors:** Sarah E. Davidson, Matthew W. Wheeler, Scott S. Auerbach, Siva Sivaganesan, Mario Medvedovic

## Abstract

Computational methods for genomic dose-response integrate dose-response modeling with bioinformatics tools to evaluate changes in molecular and cellular functions related to pathogenic processes. These methods use parametric models to describe each gene’s dose-response, but such models may not adequately capture expression changes. Additionally, current approaches do not consider gene co-expression networks. When assessing co-expression networks, one typically does not consider the dose-response relationship, resulting in ‘co-regulated’ gene sets containing genes having different dose-response patterns. To avoid these limitations, we develop an analysis pipeline called Aggregated Local Extrema Splines for High-throughput Analysis (ALOHA), which computes individual genomic dose-response functions using a flexible class Bayesian shape constrained splines and clusters gene co-regulation based upon these fits. Using splines, we reduce information loss due to parametric lack-of-fit issues, and because we cluster on dose-response relationships, we better identify co-regulation clusters for genes that have co-expressed dose-response patterns from chemical exposure. The clustered pathways can then be used to estimate a dose associated with a pre-specified biological response, i.e., the benchmark dose (BMD), and approximate a point of departure dose corresponding to minimal adverse response in the whole tissue/organism. We compare our approach to current parametric methods and our biologically enriched gene sets to cluster on normalized expression data. Using this methodology, we can more effectively extract the underlying structure leading to more cohesive estimates of gene set potency.

## Introduction

Genomic dose-response (GDR) modeling evaluates the relationship between chemical exposure and transcriptional responses to identify a level of minimal biological effect, called a point-of-departure (POD). Traditionally, PODs are estimated using *in vivo* toxicology responses. It has been proposed when minimal *in vivo* data exists, GDR data can be used in an interim risk assessment to estimate a genomic POD [1–4]. Acceptance of this approach relies on mitigating concerns about the qualitative differences between genomic and traditional PODs. A suitable estimation of transcriptional PODs accounts for differences in biological relevance between perturbed pathways and incorporates mechanistic information [2, 5].

To date, several GDR modeling approaches have been developed with a variety of methodological features, see S1 Table 1. Except for BMDExpress [6–8], which has documented guidelines in NTP et al. [9], none of the other approaches have established an approach for determining a genomic POD linked to gene- and pathway-level biological alterations. Consequently, BMDExpress [6–8] is still the most widely utilized approach for GDR modeling and will be the focus of our comparison. The platform uses a multi-step framework of parametric modeling and bioinformatics tools to characterize DR patterns and evaluate alterations in cellular function [6–9]. It addresses common challenges by using genomic data in risk assessment, but some of the assumptions made may not be appropriate given the observed data. In cases where cytotoxicity occurs in high-doses, a non-monotonic (i.e., curve with umbrella orderings) is possible [10]. For such shapes, parametric models are typically inadequate, because most of the models assume monotone changes in expression [9, 11–13], which may lead to removing genes with biological relevance due to poor model fit. Another limitation of BMDExpress is the enrichment analysis used to detect altered cellular functions and obtain genomic PODs. The enrichment analysis matches genes remaining after model fitting to a list of curated gene sets, such as the Hallmark gene sets [14–16], determined a priori. The remaining genes’ biological relevance is evaluated by their association to these sets. However, relying on curated gene sets disregards the underlying correlation structure of gene responses. These correlations may clarify understanding of mechanisms, such as co-regulation, related to developing a toxicological event.

Allocco et al. [17] demonstrated grouped genes with highly correlated expressions are likely to indicate co-regulation by a common transcription factor. Clustering-based upon similar dose-response patterns may identify co-regulation. Studies have shown Bayesian clustering methods employing Dirichlet Process mixture (DPM) models are preferred for identifying clusters of co-regulated genes, see Dahl [18], Fraley and Raferty [19], Medvedovic and Sivaganesan [20], Medvedovic et al. [21], Yeung et al. [22] among others. These methods do not always consider relationships among samples that may influence co-regulation patterns. DPM clustering methods that incorporate the relationship among samples using latent variables are shown to improve clustering, see Bar-Joseph et al. [23], Kirk et al. [24],Liu et al. [25], Lock and Dunson [26], Luan and Li [27, 28], McDowell et al. [29], Park and Kyung [30], Savage et al. [31] and references therein. However, there is not currently an approach that accounts for the underlying correlation structure of co-expressed genes with similar DR patterns.

Despite the advantage of a latent DPM model that simultaneously includes DR modeling and gene clustering, it is well known these approaches are computationally intensive. Our method, Aggregated Local Extrema Splines for High-throughput dose-response Analysis (ALOHA), first fits DR data, then uses a simpler DPM to cluster genes sharing similar DR patterns. From the DPM analysis, the method uses individual clusters to define PODs for co-regulated gene networks. The first stage of our analysis pipeline fits a flexible DR model using Bayesian local extremum splines [32], which is a shape-constrained non-parametric spline approach providing flexibility in fitting monotone, umbrella, and ‘J’ shaped responses. The second stage of our analysis inputs these fits into the context-specific infinite mixture model (CSIMM) developed by Liu et al. [25], and clusters genes with similar expression patterns. By utilizing gene fits, we incorporate GDR data’s dose-dependent nature for clustering while removing the scatter typically observed in such experiments. Biologically enriched gene sets are evaluated to obtain a POD possibly related to adverse biological responses.

ALOHA consists of two parts (i) shape-constrained non-parametric modeling on individual genes discussed in §0.1 and (ii) clustering genes with similar DR patterns covered in §0.2. In §0.3, we outline the application of our method to genomic dose-response data from HepaRG cells exposed to various compounds [33] and compare the results with the current GDR modeling and clustering approaches, respectively.

## Methods

### 0.1 Dose-Response Modeling

We observe *n* genes across *D* dose groups, with each dose group having *K* replicates such that *K* ≥ 1. For simplicity of exposition, *K* is constant across each dose group but could change at each dose level. Thus, we have *p* = *D* × *K* total observations for each gene. Let *Y* be the *n* × *p* matrix of these observations, such that the *i*^*th*^ gene is represented by the row-vector *y*_*i*_ = (*y*_(*i*,1)_,…, *y*_(*i,p*)_). The expression of gene *i* over the experimental doses can be represented as:

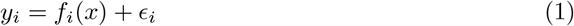

where *x* = (*x*_1_,… *x*_*p*_) is a vector of doses, *f*_*i*_ is the DR function, and ϵ_*i*_ = (_(_ϵ_*i*,1)_,…, ϵ_(*i,p*)_) is a vector of measurement errors. Here, we assume ϵ_*i*_ ∼ *N* (0, *σ*^2^*I*), where *I* is a *p* × *p* identity matrix and *σ*^2^ is some unknown variance. We use Bayesian shape-constrained non-parametric models to estimate the dose-response curve (*f*_*i*_(*x*)) for all *n* genes.

#### 0.1.1 Shape-constrained Non-parametric Modeling

For shape-constrained non-parametric modeling, we use a variant of Bayesian local extrema (BLX) splines [32]. This approach places restrictions on the shape of the model space allowing functions with no more than *H* extrema (i.e., change-points or local minima/maxima). We define the BLX spline to model functions having at most two extrema. That is, they may increase and then decrease (i.e., have an umbrella ordering), decrease and then increase (i.e., have a ‘J’ shaped ordering), or increase then decrease and increase again (i.e., have a tilde – ‘∼’ – shaped ordering). In practice, we do not consider the ‘∼’ shape plausible, but it is required to consider ‘J’ and umbrella orderings simultaneously. When a ‘∼’ shape is observed, it typically represents no dose-response relationship, and as a result, the POD for the given gene is greater than the maximum tested doses.

For gene *i*, we estimate the dose-response, *f*_*i*_, with a BLX spline defined over the fixed knot set *T* = {0, 0.5, 1}, where knots are chosen to represent the background, midpoint and maximum dose. The approach is different from Wheeler et al. [32] who consider a prior over a set of knots dense in [0, 1]. For our application, there are not enough dose groups to use this prior, and the fixed set *T* is adequate. With *H* = 2,the BLX spline model for *f*_*i*_ is:

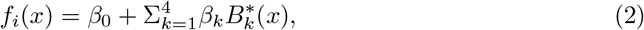

where *β*_0_ is the intercept, *B*^∗^(*x*) is a BLX spline basis function of degree 2, and *β*_*k*_ for {*k* = 1,…, 4} are spline coefficients. For more information on construction, we refer the reader to Wheeler et al. [32].

We assume dose-response relationships are initially increasing by setting the fixed integer of the BLX spline basis to one (*M* = 1). The function models monotonic increasing, monotonic decreasing, J-shaped, umbrella shaped, or ‘∼’ shaped curves (see S1 Appendix §3.1 and S1 Figs 2-7 therein). All model parameters are estimated as in [32].

Though the dose-response of genes may differ, unlike parametric modeling, the BLX spline specification is the same for each gene. This model similarity facilitates clustering on the estimated parameters (i.e., one can cluster the dose-response curve on the BLX fitted parameters). The BLX approach is different from parametric models, where the parameters all have different interpretations making clustering between models difficult.

#### 0.1.2 Parametric Modeling

For the parametric modeling approach, we estimate *f*_*i*_ using all available models, excluding the Polynomial 3 and 4, from BMDExpress (version 2.3) [6–8]. We refer the reader to Auerbach [6], Phillips et al. [7], Yang et. al. [8] for the model equations. Higher-order polynomials in most cases will fit dose-response data similarly to a quadratic model with fewer degrees of freedom and therefore were excluded from our set. This model suite captures a majority of biological response shapes thought to arise in practice. We use the model with the best fit, determined using the Aikake Information Criterion (AIC) Akaike et al. [34], Akaike [35], Sakamoto et al. [36], for inference and comparison to the BLX fit.

Other than using a quadratic dose-response curve, the parametric model suite cannot represent ‘J’ or umbrella-shaped orderings, which cytotoxic doses can induce. A quadratic model – the only parametric model that can evaluate a ‘J’-shaped response – assumes the response is approximately parabolic forcing symmetry. Symmetry around the extrema occurs infrequently. Dose-response relationships that enforce symmetry may be inadequately represented in the low-dose region.

#### 0.1.3 BLX Spline and Parametric Model Comparison

As a comparison between model fitting approaches, we compare the mean square predicted error (MSPE) between both estimates using a leave-one-out cross-validation (LOOCV) approach removing one non-zero dose group at a time, i.e.,

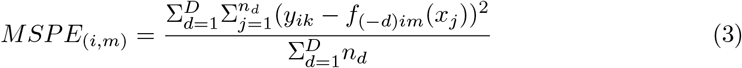

where *i* denotes the gene, *m* denotes the best parametric or BLX spline fit, *d* denotes the dose group (excluding the control group), *n*_*d*_ is the number of samples in *d, j* denotes the sample, *x*_*j*_ denotes the dose administered for sample *j*, and *f*_(−*d*)*im*_ is the model fit without dose group *d*. The MSPE measures how well each models predicts unobserved data, and it provides a comparison between each approach.

### 0.2 Gene Clustering

We identify co-expression clusters sharing similar DR patterns using the context-specific infinite mixture models (CSIMM) clustering algorithm developed by Liu et al. [25], which allows us to pre-specify sample contexts and evaluate global and local patterns. We modify this approach to model-fit data.

After dose-response modeling, the ALOHA clustering algorithm uses the BLX estimated dose-response data (e.g. red points in Fig 1) as input for the CSIMM MCMC [25]. We compare the ALOHA clustering algorithm with two other alternatives, including CSIMM and clustering on BMDExpress model-fit data. CSIMM uses the observed expression data (e.g. black points in Fig 1) and BMDExpress uses the estimated dose-response data from the best parametric models (e.g. blue points in Fig 1). All three algorithms used are given in Table 1. We compare these clustering results to evaluate their ability to capture biologically relevant co-expression clusters with coherent DR patterns and potency estimates (i.e., benchmark dose) for enriched gene sets. For further details on the CSIMM MCMC and the posterior-pairwise probability (PPP) matrix we refer the reader to §3.3 in the S1 Appendix and Liu et al. [25].

**Fig 1.**
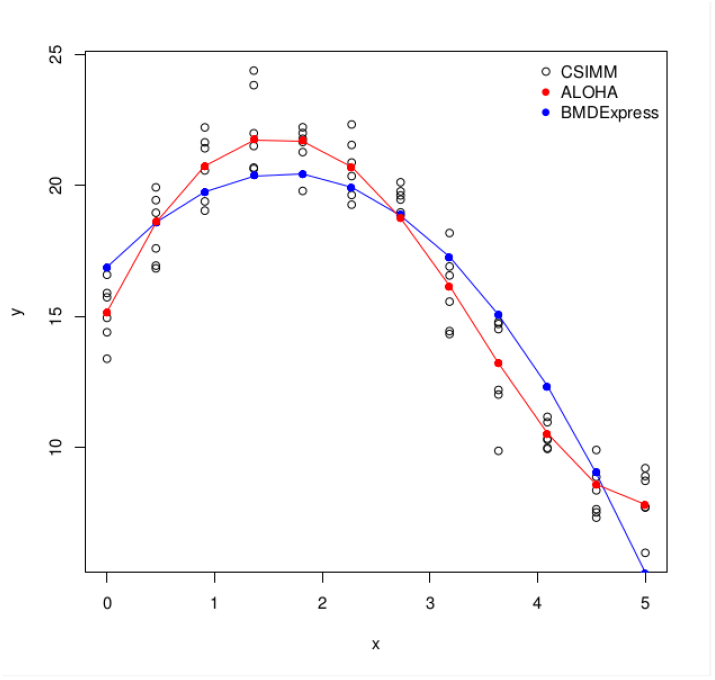
Simulated dose-response data and model fits demonstrating data used for gene clustering approaches. Black points illustrate the observed data values used as input in CSIMM. Red points illustrate the BLX estimated data values used as input in ALOHA. Blue points illustrate the estimated values from a quadratic model fit. *f* (*x*) = 7*sin*(0.3*πx*) + 15 + ϵ_*i*_, where ϵ_*i*_ ∼ *N* (0, 1).

**Table 1.** Three approaches for clustering genes to identify co-expression clusters with similar DR patterns.

| Strategy (Abbreviation) | Data | CSIMM Contexts |
| --- | --- | --- |
| Original DR Data (CSIMM) | Normalized expression data with all replicate samples | Dose groups $d \in \{1, \dots, D\}$<br>$M = D$ |
| BLX Estimated DR Data (ALOHA) | Average estimated expression for each dose group using individual BLX fits | Dose groups $d \in \{1, \dots, D\}$<br>$M = D$ |
| BPF Estimated DR Data (BMDEExpress) | Average estimated expression for each dose group using individual best parametric fits (BPF) | Dose groups $d \in \{1, \dots, D\}$<br>$M = D$ |

In addition to the clustering comparisons, we investigate the Hallmark gene sets identified in the pathway enrichment analysis used in BMDExpress. Enrichment analyses are qualitatively different, and we do not compare the enrichment scores from the two approaches. Our focus was on comparing the identified gene sets, the gene set level potency estimates (i.e., BMD and its interval) obtained from the clustering approach compared to BMDExpress, and the core genes contributing to the gene set level BMD.

### 0.3 Application

#### 0.3.1 Data

To investigate ALOHA, we use datasets on three compounds tested by Ramaiahgari et al. [33]. Of these compounds, Benzo[*α*]pyrene (BAP) and Chlorpromazine (CPZ) were the positive control chemicals, and potassium chloride (KCl) was the negative control. We evaluate the positive control compounds with and without cytotoxic doses, i.e., “all doses” (AD) and “cytotoxic doses removed” (CDR), respectively. Gene expression was measured using the S1500+ TempO-Seq platform [33, 37, 38]. All datasets’ raw RNA read counts are normalized to a log counts per million scale (*log*_2_(*CPM* + 1)). Doses were transformed to a *log*_10_(*dose* + 1) scale for approximately equally spaced dose groups. The probes that did not meet a minimum read count (MRC), i.e., median *log*_2_(*CPM* + 1) response equal to 0 and variance less than 0.1, were removed from further analysis.

#### 0.3.2 Application of BMD modeling

We fit each probe passing the MRC criteria individually as described above. For the BLX fits, a total of 50, 000 MCMC iterations were taken with the first 10, 000 disregarded as burn-in, which is are the values used in [32]. Trace plots were analyzed and showed quick convergence (usually within 200 iterations) and reasonable mixing. For parametric models, all modeling is performed with BMDExpress (version 2.3) [6–8].

Estimates of the DR are used to compute a benchmark dose (BMD) for each probe. BMD estimates for both up and down-regulation are computed by solving for *BMD* in:

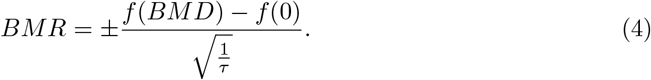

Here, *τ* ^*−*1^ is estimated variance, BMR is the benchmark response, *f* (0) is the response at the control dose [39]. In this definition, the BMR indicates a standard deviation change in gene expression from the control. For our application, we consider set *BMR* = 1, which is consistent with previous analyses with these datasets Ramaiahgari et al. [33] and NTP et al. [9].

For monotone curves, we compute the BMD using (4). Most parametric models in BMDExpress assume expression responses are monotone. In the non-monotone cases, the direction of adversity is determined using a linear trend test [6].

For BLX estimates, it is possible an up- and down-regulated BMDs exist. In these cases, we compute a BMD for both directions of adversity, where the curve’s shape determines an adverse response. For DR curves having two extrema, we consider the expression changes in the first “bump” to be an artifact of the fitting procedure. Consequently, we choose the largest BMD found after the maximum as the estimate. For DR curves that are ‘J’ or umbrella-shaped, there may be two solutions for the BMD. In these cases, the smallest of the estimates is chosen to be the BMD estimate. When no solution to (4) exists in the dosing interval, we set the BMD to be one plus the maximum experimental dose (i.e. *x*_*D*_ + 1) to place the BMD above the maximum experimental dose. If up- and down-regulated BMDs exist, we use the smallest of these BMDs as the estimate.

Confidence limits in the BMD are estimated relative to the estimation method. For the BLX spline, calculated using MCMC, we take the 2.5^*th*^ and 97.5^*th*^ quantiles of the BMD distribution to estimate the BMDL and BMDU, respectively. For parametric models, estimated using maximum likelihood, the limits are estimated using the profile likelihood. Point and interval estimates of each BMD are transformed from the log-scale to the original dose scale before calculating the ratio of the upper and lower confidence limits (BMDU/BMDL). The BMDU/BMDL ratio is a measure of reliability in the BMD estimate of each probe. Smaller BMDU/BMDL estimates indicate increased certainty in BMD estimates and more reliable fits. We define probes with reliable expression activity as those having a (i) BMD less than the maximum dose and (ii) 95% confidence limit BMDU/BMDL less than 40, which are consistent with the reliability criteria proposed by NTP et al. [9]. When fitting parametric models, the best model fits must also have a goodness-of-fit p-value *>* 0.1, which indicates no significant difference between the observed expression values and those estimated by the model. Probes meeting this criterion have reliable expression activity and remain in the analysis. This is consistent with NTP et al. [9].

#### 0.3.3 Gene Clustering

After model fitting, model data from reliable probes are used to obtain the inputs, described in §0.2, to the CSIMM clustering algorithm.

In the TempOSeq platform, multiple probes target one gene designated by a common EntrezID, but their targets may be different gene isoforms displaying different co-expression patterns. Consequently, we do not average these probes’ responses, and we remove probes without a designated EntrezID or multiple EntrezIDs.

For CSIMM, the MCMC sampler is run for 50,000 iterations, with the first 10,000 samples disregarded as burn-in. The clustering strategies are assessed by their ability to separate DR trends into co-expression clusters, where separation is assessed using a heat map. Biological enrichment of clustering results is analyzed using Cluster Enrichment Analysis (CLEAN) [40]. This approach compares the level of functional coherence in clusters based on previously curated biological pathway lists and potency estimates for enriched gene sets.

To compute CLEAN scores, we use the human Hallmark Gene List and the Computational Cancer Gene Neighborhood Gene List (C4:CGN or CGN) from the Molecular Signatures Database (MSigDB) to evaluate the enrichment of gene clusters [14–16, 41]. Gene sets containing at least three genes are used for the enrichment analyses consistent with NTP et al. [9]. A gene set is enriched for a cluster if Fischer’s exact test Benjamini-Hochberg False Discovery Rate adjusted p-value (FFDR) is less than 0.01. We consider gene clusters with the smallest FFDR p-value for at least one enriched gene set to be the most relevant. For each enriched gene set to be considered for biological interpretation, based on the criteria from NTP et al. [9], the most relevant cluster representing it must contain at the minimum 3 of its genes and populate the set by at least 5%. Though we include the NTP et al. [9] criteria for defining active pathways since we include the FFDR p-value threshold criteria for the clustering approaches, our criteria are fundamentally different from those used in Phillips et al. [7], Yang et al. [8], NTP et al. [9].

Potency estimates for enriched gene sets are obtained using the average BMD estimates from the core genes representing each enriched gene set. We define the core genes to be those in the most relevant cluster for an enriched gene set belonging to that set. For parametric fits, potency estimates are the median BMD estimate of the enriched gene sets. For the clustering results on the normalized expression data and the BLX spline fits we use the posterior mean BMD estimate for each gene. We obtain the lower and upper bounds on the gene set BMD – BMDL and BMDU, respectively – using the 0.025 and the 0.975 quantiles of the best BMD estimates for core genes to obtain a 95% credible interval.

#### 0.3.4 BMDExpress Functional Classification Analysis

After parametric model fitting and post-model filtering of probes, we performed functional classification analyses in BMDExpress using the human Hallmark Gene List from the Molecular Signatures Database (MSigDB) to compare gene set level BMD estimates with those from each of the clustering approaches [6–8, 14–16, 41]. It should be noted, prior to the functional classification analysis, BMDExpress consolidates multiple probes mapping to the same EntrezID. The BMD of these probes are averaged together to obtain one BMD value representing their common EntrezID [6–8].

From the pathway enrichment results from BMDExpress we obtain a gene set level potency estimate for each of the gene sets. For our analysis we use the median BMDL, median BMD, and median BMDL as estimates of the potency estimates for each of these gene sets. We compare these estimates with those obtained from the most biologically relevant cluster for each of the enriched Hallmark gene sets found with various clustering approaches.

## Results

### 0.4 Model Fitting

#### 0.4.1 Model Predictions

We evaluate each modeling approach’s ability to obtain the best estimate of the dose-response curve using a leave-one-out cross-validation (LOOCV) prediction. Fig 2 shows the mean squared prediction error (MSPE) for the parametric and BLX fits for each probe attaining the minimum response criteria in §0.3.1 and having less than 75% of their observed responses equal to zero. A smaller MSPE indicates a better prediction. In this figure, the blue dashed line shows situations where the MSPE is the same for a given probe. Black dots, or points to the right of the line, show results where ALOHA outperforms the parametric approach. Red dots, or points to the left of the line, show results where the best parametric model is superior. Here, most of the red points are close to the blue line, which indicates the method fit and predictions are essentially the same for that probe; however, the black points show a greater right skew. ALOHA does a better job describing the dose-responses for these probes, which occurs for approximately 50% of the probes. Further, we observe in general the maximum of the BLX spline MSPEs is an order of magnitude smaller than the maximum of the MSPEs for the best parametric fit, which gives credence to the argument that ALOHA is better representing the true underlying dose-response curve.

**Fig 2.**
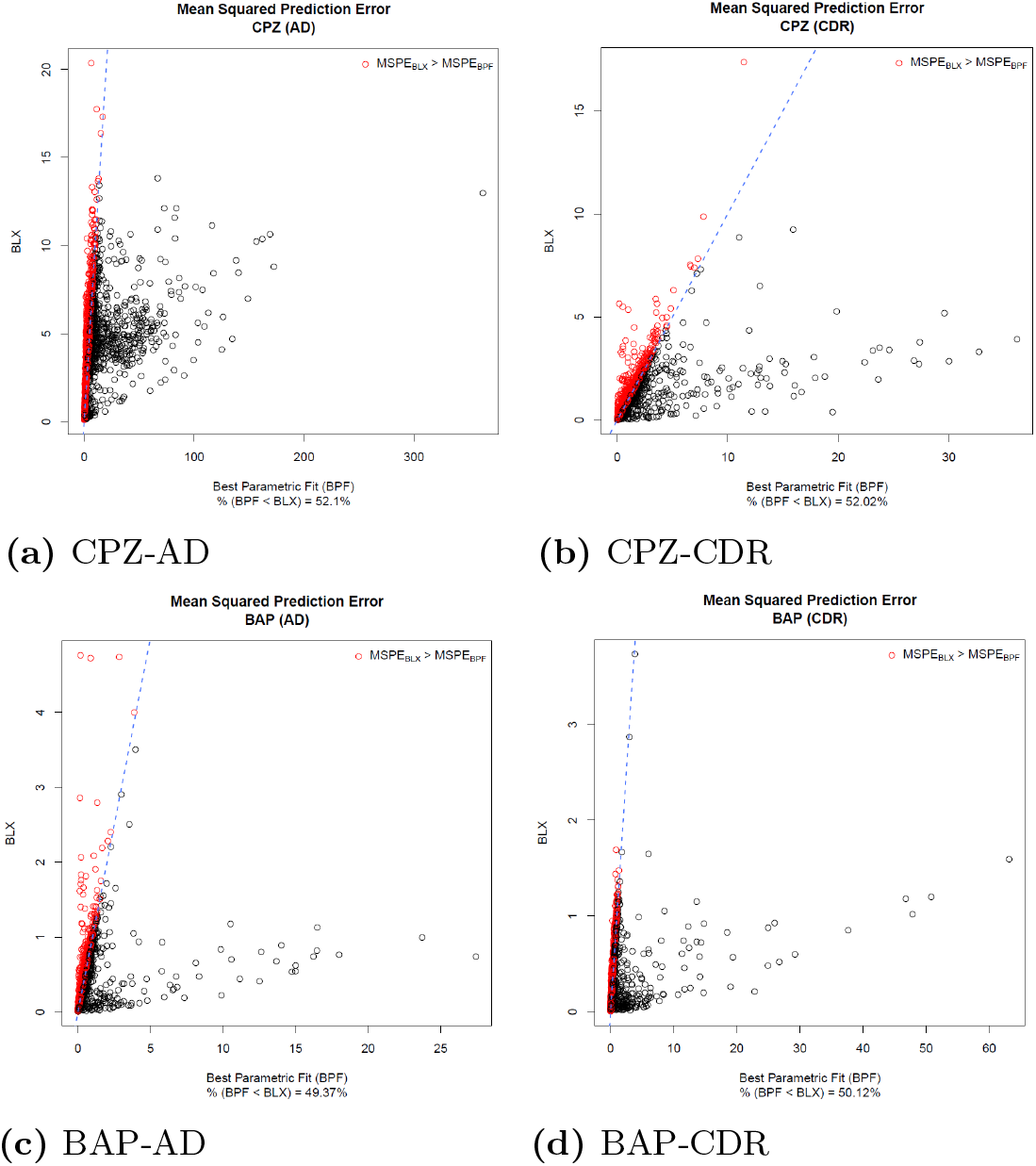
Scatterplots with the mean squared prediction error (MSPE) for the best parametric model fit (BPF) (x-axis) and the BLX spline (y-axis) for each MRC filtered probe modeled with both methods and having less than 75% percent of observations with a response equal to zero. The blue dashed line indicates where the BLX and BPF MSPE are equivalent. Red points indicate probes with a smaller BPF MSPE than the BLX MSPE. Black points indicate probes with a BLX MSPE smaller than or equal to the BPF MSPE.

Fig 3 shows why ALOHA is doing better modeling these data and compares the curve fit for a probe from Benzo-[*α*]-pyrene. The left figure has all doses and while the right one has the highest cytotoxic dose removed. In the left plot, we observe a clear non-monotonic response which the BLX spline fit captures; however, the Exponential 5 model misses this behavior. The right plot shows the fits where the highest dose group is removed since it is cytotoxic. Though this dose’s removal is to mitigate issues of non-monotone relationships, there is still a roughly non-monotonic shape. In both of these cases, the model constraints of the best parametric fit, i.e. the Exponential 5, induces a plateau in the high doses, which is unable to capture the peak response and the change in response resulting in a poor fit to the observed data (i.e. fit p-value = 1*e* − 4 for both Benzo-[*α*]-pyrene with and without cytotoxic doses). On the other hand, our approach’s flexibility captures the non-monotonic response when all doses are included, and it splits the difference between the response in the two highest dose groups when cytotoxic doses are excluded. There is no longer sufficient evidence to determine if there is a true non-monotonic response.

**Fig 3.**
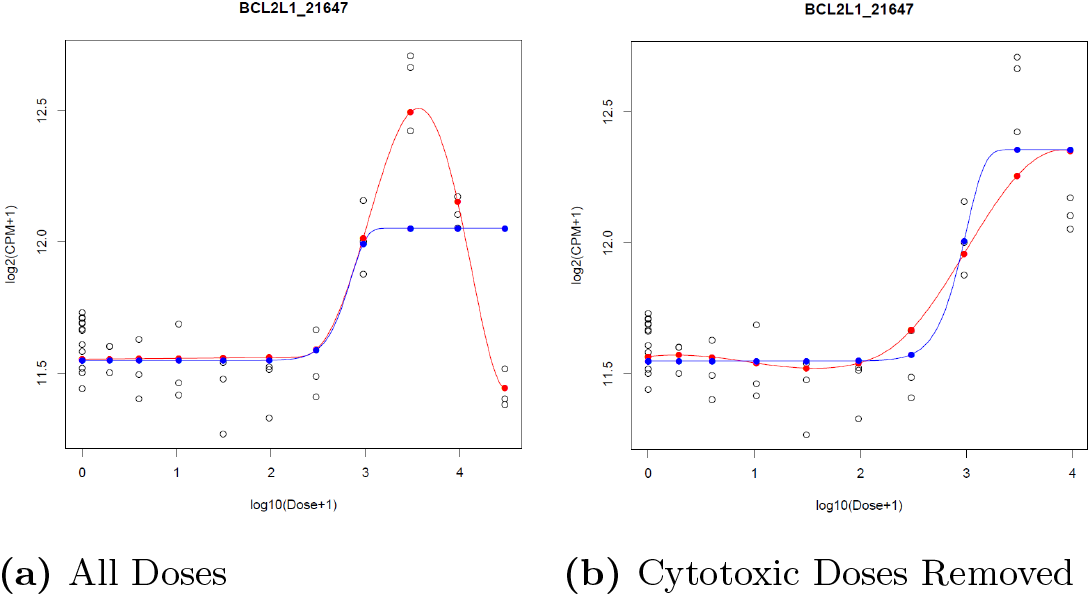
Probe passing the reliability criteria with BLX spline model fitting but fails with BMDExpress (version 2.3) parametric modeling for Benzo-[*α*]-pyrene all doses (left plot) and cytotoxic doses removed (right plot). In both plots the curves display the BLX spline (red) and best parametric model fit from BMDExpress (blue).

#### 0.4.2 Probe Retention

Out of 2977 probes in the S1500+ TempO-Seq platform, there were 2925, 2703, 2684, 2928, and 2922 probes used in the analysis for Potassium chloride, Benzo-[*α*]-pyrene with and without cytotoxic doses, and Chlorpromazine with and without cytotoxic doses, respectively, based on MRC filtering described in §0.3.1. Table 2 shows the number of modeled probes passing the criteria described in §0.3.2 with the two modeling approaches. Other than Benzo-[*α*]-pyrene with all doses and Chlorpromazine without cytotoxic doses, more probes passed with parametric modeling compared to BLX spline modeling. Many of the same probes were retained between the two modeling approaches.

**Table 2.**
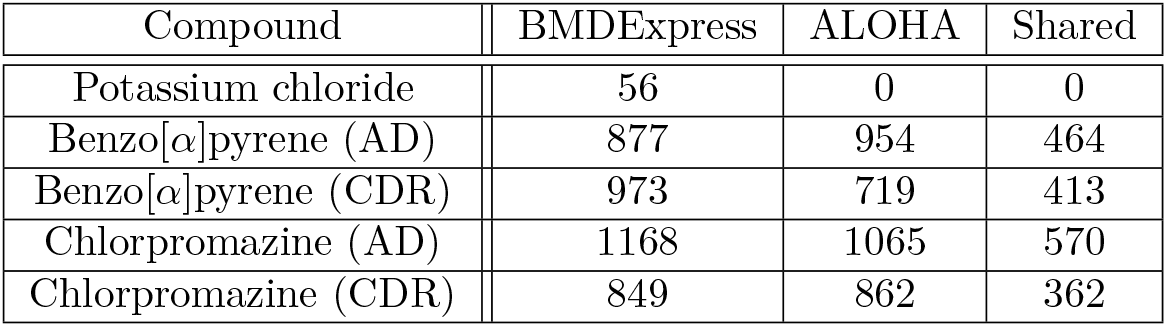
Number of modeled probes passing model-fit criteria in BMDExpress (parametric models) and ALOHA (BLX spline model), respectively. Columns 2 & 3 provide the number of probes passing the criteria in §0.3.2 for each dataset. Column 4 provides the number of probes in common between the modeling approaches. [AD - all doses; CDR - cytotoxic doses removed].

| Compound | BMDEExpress | ALOHA | Shared |
| --- | --- | --- | --- |
| Potassium chloride | 56 | 0 | 0 |
| Benzo[$\alpha$]pyrene (AD) | 877 | 954 | 464 |
| Benzo[$\alpha$]pyrene (CDR) | 973 | 719 | 413 |
| Chlorpromazine (AD) | 1168 | 1065 | 570 |
| Chlorpromazine (CDR) | 849 | 862 | 362 |

### 0.5 Gene Clustering Comparison

#### 0.5.1 Separation of Dose-Response Curve Shapes

For both Benzo-[*α*]-pyrene and Chlorpromazine, normalized expression values for up and down-regulated genes were better separated with the CSIMM (see S1 Figs 19-22). In this case, the BLX spline parameter estimates showed poor separation, which implies dissimilarity in dose-response curves for genes in the same cluster. For the ALOHA approach, the BLX parameter estimates were separated, which indicates greater similarity in the shape of dose-response curves for genes belonging to the same.

Separation of BLX estimated dose-response data were also evaluated to confirm the coherence of dose-response trends of genes in the same cluster. We chose to cut the posterior pairwise dendrogram such that five representative clusters resulted. The chosen cut-off was arbitrary, but was done for exposition of dose-response trend coherence within clusters. For both Benzo-[*α*]-pyrene and Chlorpromazine, the ALOHA clusters had a high level of coherence in the dose-response patterns compared to CSIMM (see Fig 4 & S1 Figs 23-26). We also calculated several internal validation scores for the five resulting clusters using the Euclidean distance as the measure of dissimilarity [42–45] (see Table 3 & S1 Tables 2-5). All validity scores were consistent with greater coherence in dose-response trends of ALOHA gene clusters compared with CSIMM. These results indicate ALOHA is better suited for identifying gene groups with co-regulated dose-response patterns.

**Fig 4.**
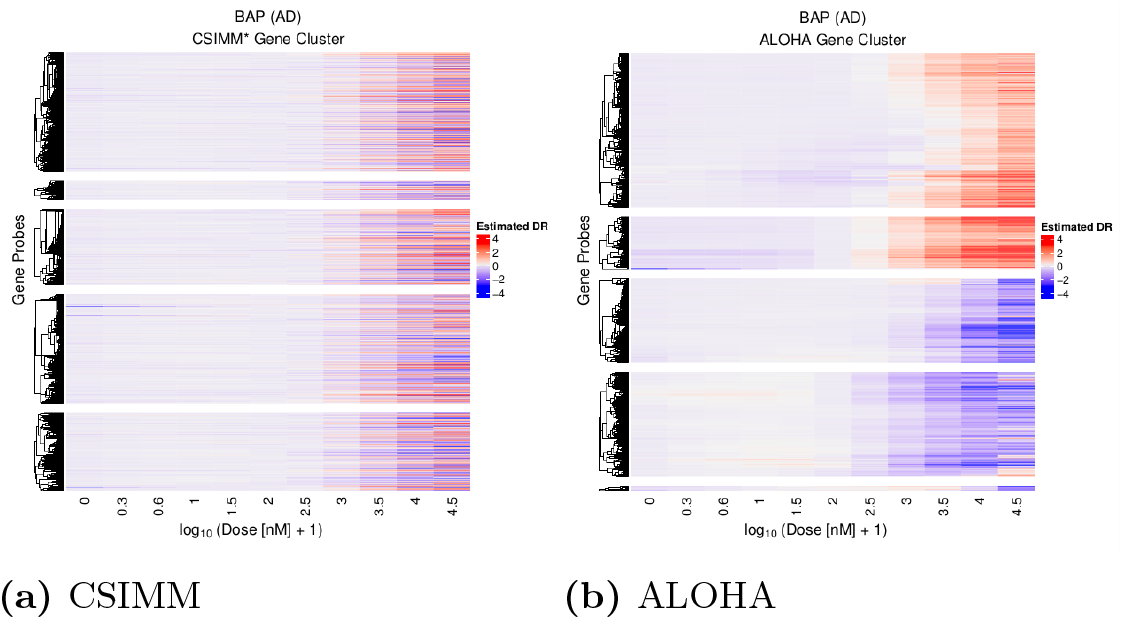
BAP AD - Heatmaps displaying clustering results applied to BLX estimated dose-response data showing visual separation of DR trends. (Left - *CSIMM*) CSIMM clustering on observed normalized expression data with all replicates in each dose group. (Right - *ALOHA*) CSIMM clustering on BLX estimated dose-response data.

**Table 3.**
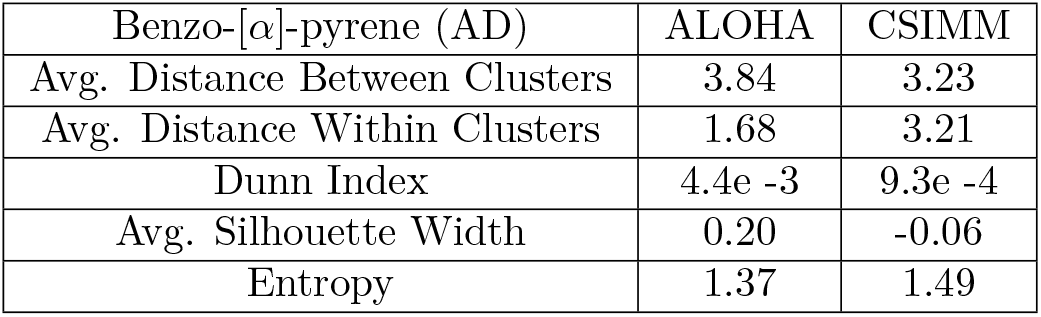
Scores measuring the internal coherence of dose-response trends in the five clusters resulting from ALOHA and CSIMM in the Benzo-[*α*]-pyrene dataset with all doses, see Fig 4. Larger values for average distance between clusters, Dunn index, and the average silhouette width indicate better separation of genes into similar dose-response groups. Smaller values for the average distance within clusters and entropy indicate more coherence between genes clustered together.

#### 0.5.2 Functional Coherence in Clusters

CLEAN scores [40] were used to compare the quality of functional coherence in the resulting co-expression clusters. Fig 5 compares the empirical cumulative distribution function (ECDF) of CLEAN scores for all clustering approaches on Benzo-[*α*]-pyrene with all doses. For BMDExpress clustering, there were fewer genes with a non-zero CLEAN score. Further, the CLEAN scores were generally smaller, e.g., the Benzo-[*α*]-pyrene with all doses results showed no biological significance with BMDExpress clustering. Generally, BMDExpress was the worst of the three approaches at identifying biologically relevant clusters of the three approaches (see S1 Figs 27-30). There was little separation between the ECDF curves for CSIMM (black) and ALOHA (red), except in the case of Chlorpromazine with all doses. In that case, there was no clear delineation between any of the curves, which indicates that both approaches obtained similar levels of functional coherence in co-expression clusters (see S1 Figs 27-30). Enrichment scores from the most relevant cluster of the enriched gene set showed biological relevance of the clustering approaches was similar (see S1 Figs 31-38). Although CSIMM and ALOHA had similar functional coherence, as seen below, CSIMM did not produce clear co-regulated dose-response functions, i.e., the dose responses for genes in a given cluster were not similar (see Fig 4).

**Fig 5.**
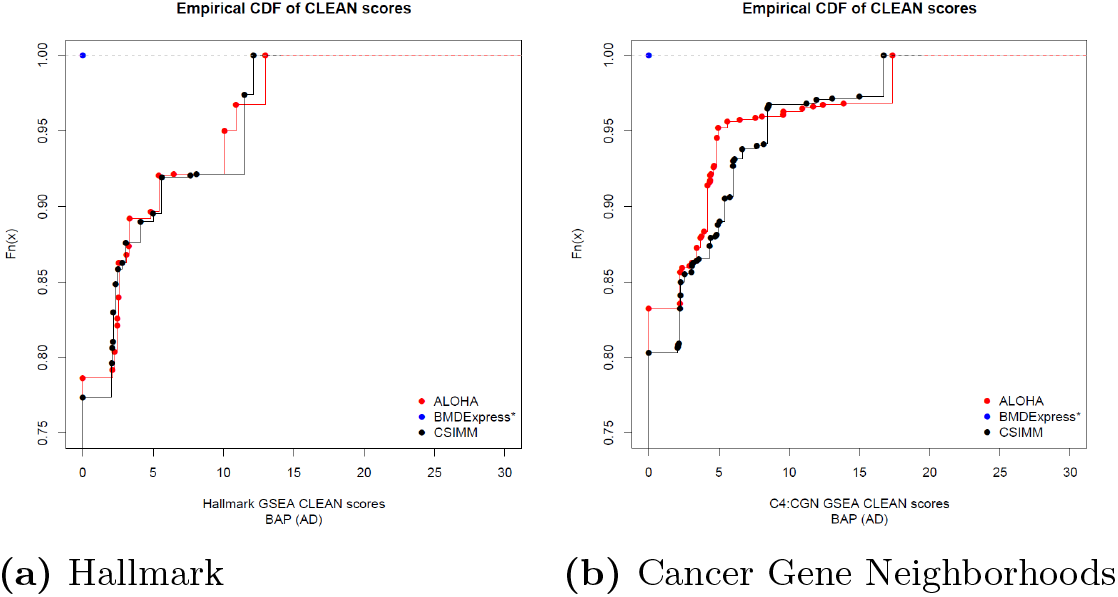
CLEAN score empirical cumulative distribution function (ECDF) plots for Benzo-[*α*]-pyrene including all doses (BAP-AD), with MSigDB’s Hallmark (right) and Cancer Gene Neighborhood (CGN) (left) gene lists on the normalized expression (CSIMM – black), BLX estimated expression (ALOHA – red), and best parametric fit estimated expression (BMDExpress – blue) approaches.

#### 0.5.3 Benchmark Dose (BMD) Coherence

Fig 6 shows the BMD estimate, and the associated 95% credible interval estimate for each enriched Hallmark gene set from the clustering approaches. For enriched gene sets shared between CSIMM and ALOHA, the gene set level BMD estimates were similar. There were a few exceptions to this behavior. For example, in gene set “e2f_targets”, the ALOHA cluster BMD interval was smaller, indicating improved BMD coherence. Generally, gene set BMD intervals for BMDExpress clustering were wider than those from ALOHA.

**Fig 6.**
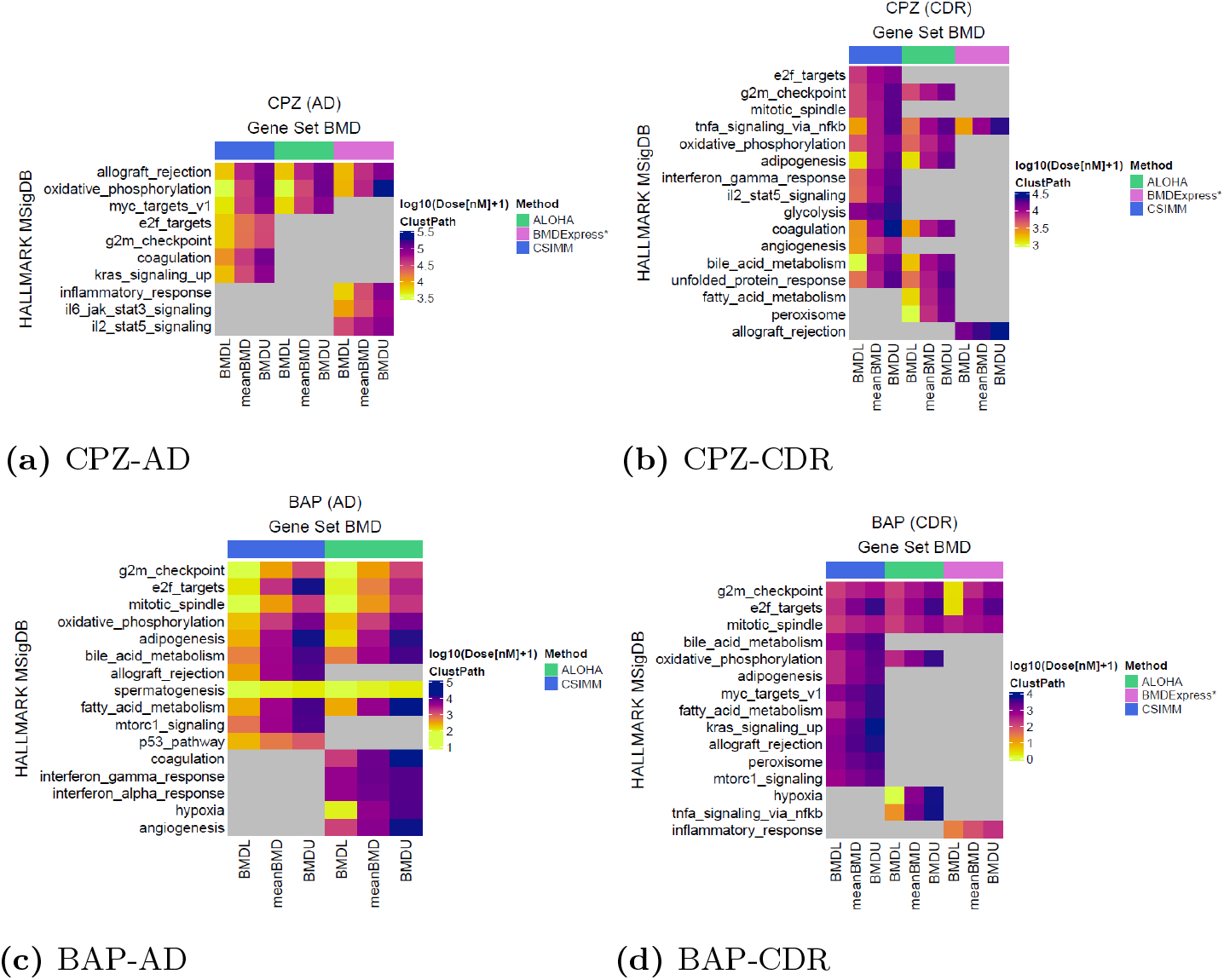
Heatmaps comparing the posterior mean BMD and 95% credible intervals (i.e. BMDL and BMDU) estimates (columns) for enriched Hallmark gene sets [14–16] (rows) across the three clustering methods. The BMDs are estimated using core genes in the most relevant cluster, as defined in §0.3.3, mapping to the respective gene sets. (a) Chlorpromazine with all experimental dose groups included, (b) Chlorpromazine with cytotoxic dose groups removed, (c) Benzo-[*α*]-pyrene with all experimental dose groups, (d) Benzo-[*α*]-pyrene with cytotoxic dose groups removed. Gray blocks within the heatmap indicate which gene sets are not enriched for a clustering approach. Bars on above the columns indicate the various clustering approaches, i.e. CSIMM (blue), ALOHA (green), and BMDExpress clustering (magenta, described in §0.2.

Similar trends were observed for the enriched Cancer Gene Neighborhood gene sets shared between the clustering approaches, except for the Chlorpromazine with all doses which had tighter BMD intervals for many of the gene sets enriched with BMDExpress clustering (see S1 Figs 39-41). Though there were similar BMD estimates between CSIMM and ALOHA, the gene clusters from CSIMM were not representative of a particular dose-response pattern. As a result, BMD estimates were inconsistent about the direction of regulation, and they provided contradictory information about the adverse direction.

Gene set BMD estimates from the clustering approaches were also compared with those estimated after functional classification in BMDExpress described in §0.3.4. Overall, these comparisons showed more genes contributed to the BMD estimate from the current BMDExpress approach (see Fig 7). With ALOHA and the other clustering approaches, individual gene BMDs were tighter grouped than those from the current BMDExpress approach. Confidence intervals for gene set BMDs with the clustering approaches also overlapped or contained those intervals from the current BMDExpress approach, which indicated clustering resulted in gene groups that produced BMD estimates consistent with BMDExpress.

**Fig 7.**
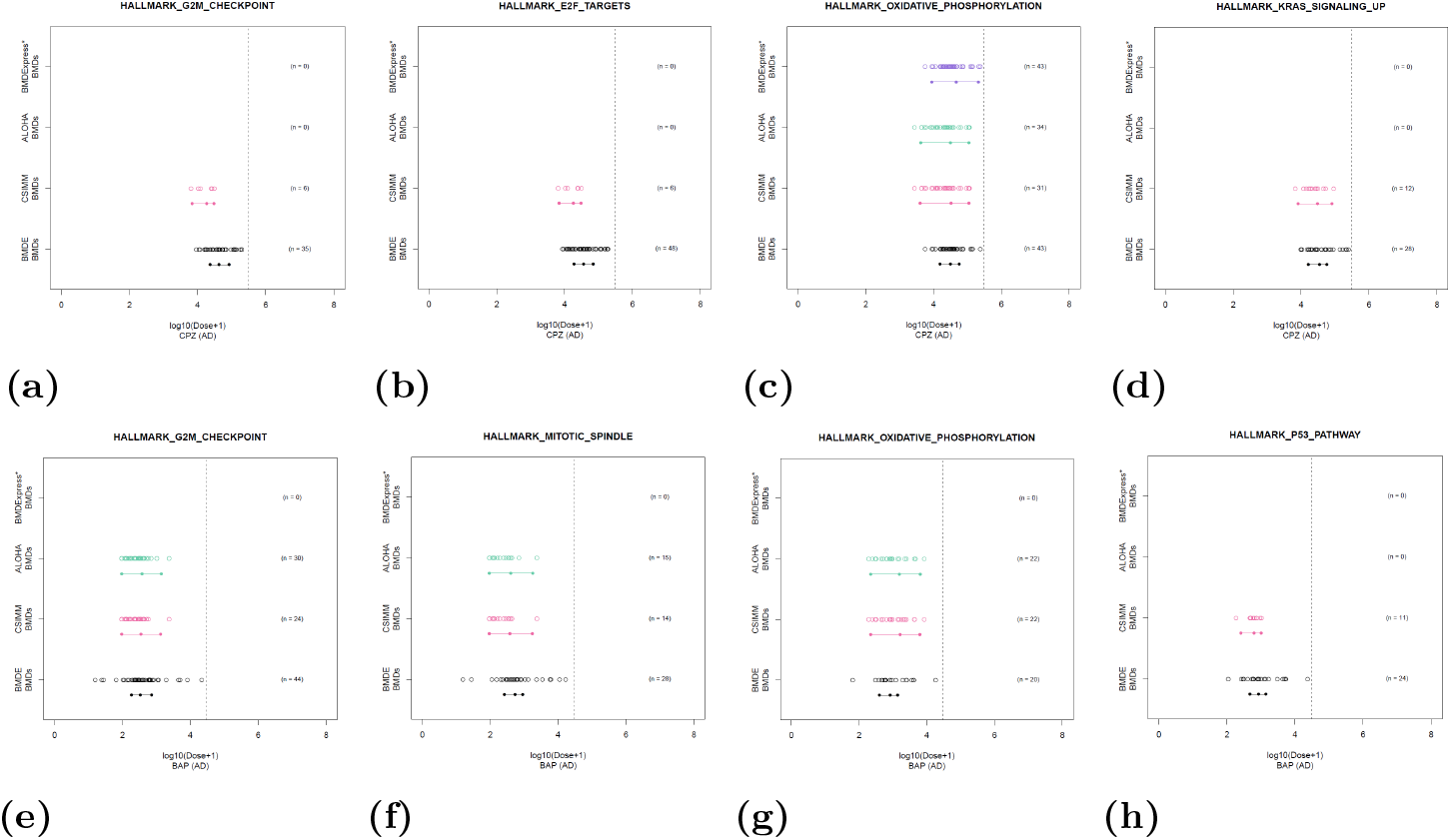
Gene set level BMD intervals from BMDExpress (BMDE), using functional classification without clustering, and the clustering approaches on Chlorpromazine and Benzo-[*α*]-pyrene with all doses (top and bottom, respectively). Points show the best BMD estimate of individual core probes and the number of core probes displayed on the right (see S1 Tables 6-13 for details about these probes). (Note: For BMDExpress, probes mapping to the same EntrezID had their BMD estimates averaged and share a single BMD value in the results, but we consider these probes individually in the count to be consistent with core probe counts with clustering.). The vertical dashed line indicate the maximum experimental dose and the line segments below the individual BMDs display the interval (i.e. BMDL to BMDU) and central tendency for the gene set level BMD as described in §0.3.3 & §0.3.4.

## Discussion

The use of genomic data in risk assessment relies on developing new methods that account for biological phenomena affecting the underlying data structure. Here, we developed a genomic dose-response (GDR) modeling approach, called Aggregated Local Extrema Splines for High-throughput Dose-Response Analysis (ALOHA), that utilizes model-fit data from Bayesian Local Extremum (BLX) splines and context-specific infinite mixture models (CSIMM) to obtain gene clusters with similar dose-response curves.

For ALOHA, the increased number of probes with a minimum read count (MRC) filtering did not impact the number of probes deemed to have dose-response (DR) information. For example, few probes were removed in the Potassium chloride dataset before fitting, but after DR modeling, there were no remaining probes (see Table 2). This result corresponds to Potassium chloride being a negative control. For probes passing only with parametric modeling, extreme response observations impacted the fit, resulting in an unrealistic fit to the data but an acceptable benchmark dose (BMD) estimate. Here, these observations did not impact the BLX fit, which resulted in greater uncertainty in the BMD estimate or a flat dose-response curve. For probes passing only with BLX, the model flexibility captured non-monotone responses in high doses where cytotoxicity may occur (see Fig 3). In these cases, parametric models were unable to fit these observations due to model constraints, resulting in poor fit estimates.

Comparing our method’s ability to describe the genomic dose-response data with BMDExpress (version 2.3), using a leave-one-out cross-validation (LOOCV) method, demonstrated little difference between the two approaches in the case of monotone dose-response curves. However, in cases where cytotoxicity induced a non-monotonic response, ALOHA produced smaller mean squared predicted error (MSPE) estimates, thus outperforming the predictive ability of BMDExpress (see Fig 2). For many of these genes, the parametric models were unable to appropriately capture the observed response at high doses due to the assumption of monotonicity and model constraints. This assumption led to poor model fits and removal from the analysis. When cytotoxicity is present, a common approach to prevent discarding genes from the analysis is to remove the doses producing the cytotoxic behavior. Though this may resolve some of the genes with non-monotone behavior, it does not guarantee this across all genes. Fig 3 illustrates this case, where removing the high doses in the Benzo-[*α*]-pyrene data set did not remove the downturn for “BCL2L1_21647”. Since the identification of cytotoxic dose groups is an arbitrary decision made by the modeler, any removal approach may result in responsive genes being arbitrarily removed from the analysis. ALOHA does not suffer from this difficulty and retains non-monotone dose-responses for further analyses.

Except for Benzo-[*α*]-pyrene and Chlorpromazine, with and without cytotoxic doses respectively, more probes passed the post-model fit criteria with BMDExpress compared to ALOHA. Investigation of individual probe fits showed BMDExpres inappropriately identified a portion of these as active (i.e., distinct and reliable DR trend). Here, probes fell into one of two categories. First, observed responses were noisy with a few extreme observations (see S1 Fig 10). Second, the response in high doses was split between active and dead cells (i.e. zero expression) – (see S1 Fig 11). In the first case, ALOHA estimated a flat curve. In the second case, ALOHA split the difference between active and dead samples’ responses in each dose group. The cytotoxic dose range uncertainty did not provide enough evidence to classify the gene as active; either the curve was flat, or there was uncertainty in the BMD estimate near the maximum experimental dose. In both of these cases, constraints in the parametric models were influenced by the most extreme response values from sensitive samples, which produced a curve resulting in a BMD estimate (see examples in S1 Figs 10 & 11). Whereas ALOHA was robust to sensitive samples, and the overall fit was unaffected without more evidence of an extreme change in response.

ALOHA’s ability to filter active genes is shown with potassium chloride, a negative control compound. ALOHA removed all probes from the analysis, unlike BMDExpress, which retained several genes as active. Some probes in the positive control compound datasets, i.e. Benzo-[*α*]-pyrene and Chlorpromazine, passed the model-fit criteria with BMDExpress but not with ALOHA. Here, the posterior distribution of BMD estimates from ALOHA was either too uncertain or bi-modal with one-mode larger than the maximum dose. In these cases, the 95% credible interval or the posterior mean BMD estimate was unable to pass the recommended criteria presented in the NTP guidance document [9]. Even without differential gene expression (DEG) filtering and increased data, ALOHA retained active probes (i.e. probes with distinct and reliable DR trends) while removing inactive ones.

Assessing ALOHA’s clustering behavior, we first compared the separation of dose-response patterns in the normalized expression responses and BLX spline parameter estimates – a proxy measure of model fit shapes – between ALOHA and CSIMM [25] (see S1 Figs 19-22). ALOHA showed no clear separation in the regulation trends of normalized expression data. However, there was a clear improvement in the separation of the BLX parameter estimates, which implies DR patterns of individuals’ genes are more coherent in ALOHA clusters than CSIMM clusters. To verify these results we compared the separation of the BLX estimated dose-response data, which confirmed the ALOHA clusters contained genes with more coherent DR patterns (see Fig 4 or S1 Figs 23 & 26).

Comparison of the functional coherence of the resulting gene clusters demonstrated ALOHA performed similarly to CSIMM. CSIMM did obtain more enriched gene sets than ALOHA. However, the clusters enriched for these gene sets were not necessarily reflective of genes with common DR patterns. Members of a gene set may have different DR curve shapes. Consequently, ALOHA breaks these sets into different clusters, whereas CSIMM clusters these genes regardless if they have conflicting DR patterns. ALOHA also showed clear improvements in the biological relevance of clusters obtained compared to BMDExpress clustering. These results were likely the result of qualitative differences in probes passed to the clustering process. Alternatively, the limited flexibility of parametric models and inactive probes’ retention may have made it difficult for CSIMM to discern clear DR co-expression clusters using BMDExpress model fit data. In these cases, additional information is needed to distinguish DR patterns better.

Gene set BMD estimates from ALOHA clustering were tighter in most cases than those from BMDExpress clustering. The only exception was for the Cancer Gene Neighborhood (CGN) gene sets enriched in Chlorpromazine with all doses. This improvement was due to clustering probe fits with more similar DR curves producing consistent BMD estimates. BMD intervals from ALOHA also largely overlapped with intervals estimated using the functional classification results from BMDExpress, which places all probes passing the model-fit criteria into their respective gene sets. Though BMDExpress generally produced tighter BMD intervals than ALOHA, the qualitative difference between the two approaches when estimating the confidence intervals was likely the source of this disparity in precision. This result is supported by the observed differences in interval estimates between BMDExpress clustering and the current BMDExpress approach. Because both use functional classification, they should be relatively similar (see “OXIDATIVE_PHOSPHORYLATION” in see Fig 7). Individual gene BMDs that contributed to the overall gene set BMD were also more diffuse than ALOHA when using the current BMDExpress approach. This result shows that even with the addition of clustering, ALOHA was able to obtain BMD estimates consistent with the current approach, BMDExpress, while identifying gene groups with more coherent BMD estimates.

Genomic dose-response (GDR) analyses currently separate modeling dose-dependent gene expression changes from characterizing coordination among responses. ALOHA integrates these analyses to identify co-expressed genes and characterize altered cellular processes. Our study evaluated three chemical compounds, two eliciting strong transcriptional responses and one with limited to no biological response. Here, ALOHA demonstrated better characterization of GDR relationships, particularly for genes with a non-monotonic response. It did this by retaining more biologically active genes while removing inactive or noisy probes by utilizing shape-constrained splines. The coupling of flexible modeling with co-expression clustering resulted in biologically relevant clusters and potency estimates consistent with the current approaches in BMDExpress.

ALOHA stream-lines current methods and better accounts for the underlying data structure. Identifying gene clusters with coordinated DR expressions captures cellular processes related to adverse health effects from which gene set level BMDs can be estimated. These estimates may be used to determine a genomic POD comparable to those derived from *in vivo* toxicological endpoints. In summary, ALOHA provides an alternative step-wise method integrating improved DR models as well as consideration of coordinated expression changes to obtain BMDs that may be used to estimate a biologically relevant POD.

While we have demonstrated ALOHA is competitive with the current BMDExpress, there are several areas for future research. The first is to evaluate the sensitivity and specificity of ALOHA’s GDR modeling strategy by including simulated dose-response data. No simulation method to generate this kind of data exists, and developing such a simulation methodology to evaluate ALOHA is an important area of future research. The second would be to optimize the clustering algorithm to obtain disjoint gene clusters that provide a clear delineation for evaluating the accuracy of similar DR patterns in gene clusters and their related biological mechanisms. The last would be to improve the estimation of uncertainty in BMDs from individual and gene set clusters. Such a refinement would allow ALOHA to better quantify the uncertainty between transcriptional and *in vivo* toxicological points-of-departure.

## Acknowledgments

We would like to acknowledge Ms. Davidson’s mentor Dr. Eileen D. Kuempel for her guidance and input on investigating the use of toxicogenomics data in occupational health risk assessment, Mr. Nathan M. Drew for his comments on technical issues during initial methods development, Mr. Wen Niu for uploading cleaned expression sets into the iLincs database, the CDC OAMD SCBS team for assistance with improving methods for high-performance computing during pipeline development, and thank Drs. Pierre Bushel and Fred Parham for reviewing the manuscript and their comments.

This work has been supported by the contract from The National Institute for Occupational Safety and Health (75D30118P01991), the grants from National Institutes of Health (U54HL127624 and P30ES006096), in part by intermural funds at National Institute of Environmental Health Sciences Division of Intramural Research (NIEHS/DIR), and in part by an appointment to the Research Participation Program at the Centers for Disease Control and Prevention administered by the Oak Ridge Institute for Science and Education through an interagency agreement between the U.S. Department of Energy and the Centers for Disease Control and Prevention.

## 1 Contributions

Sarah E. Davidson performed methods development, software development, model implementation, evaluation of results, and visualizations.

Matt Wheeler provided input on implementation and interpretation of BLX spline modeling, as well as interpretation of results.

Scott Auerbach provided experimental data, BMDExpress analysis results, and toxicological input on modeling results.

Siva Sivaganesan suggested and provided input on the model fitting leave-one-out cross-validation analysis.

Mario Medvedovic provided input on implementation, interpretation, and visualization of clustering results.

## 2 Abbreviations

ALOHA: Aggregated LOcal extrema splines for High-throughput Analysis
AD: All Doses
AIC: Aikake Information Criterion
BAP: Benzo-[*α*]-pyrene
BLX: Bayesian Local Extrema
BMD: Benchmark Dose
BMDL: lower BMD uncertainty bound
BMDU: upper BMD uncertainty bound
BMR: Benchmark Response
BPF: Best Parametric Fit
CDR: Cytotoxic Doses Removed
CGN: Cancer Gene Neighborhoods
CLEAN: Cluster Enrichment Analysis
CPM: Count per Million
CPZ: Chlorpromazine
CSIMM: context-specific mixture models
DR: dose-response
DPM: Dirchlet Proces Mixture Model
ECDF: Empirical Cumulative Distribution Function
FFDR: Fischer’s exact test Benjamini-Hochberg False Discovery Rate adjusted p-value
GDR: genomic dose-response
KCl: Potassium chloride
LOOCV: leave-one-out cross-validation
MCMC: Markov Chain Monte Carlo
MSigDB: Molecular Signature Database
MSPE: Mean squared predicted error
MRC: Minimum Read Count
POD: point-of-departure

## Supporting information

**S1 Appendix. Supplemental Material**. The appendix contains supplemental figures, tables, and equations for the reader.

**S1 Table 1. Available Genomic Dose-Response Methods Comparison Table**.

**S1 Fig 1. Comparison Outline Supplement Diagram**.

**S1 Figs 2-7. BLX Spline Curve Shapes (Extrema = 2 & Increasing = TRUE)**.

**S1 Figs 8 & 9. BMD Estimation BLX Spline Modeling**.

**S1 Clustering. CSIMM Gene Clustering Information**.

**S1 Figs 10-13. Fit Comparison Cases**. These plots provide fit comparisons in cases where the parametric fit passes but the BLX spline fails to pass.

**S1 Figs 14-18. BMD Estimate Comparison**.

**S1 Figs 19-22. Heatmaps of Dose-Response Clustering Comparisons - Observed Expression Data**.

**S1 Figs 23-26. Heatmaps of Dose-Response Clustering Comparisons - Estimated Dose-Response Data**.

**S1 Tables 2-5. Internal Clustering Validation Score Comparison Tables**.

**S1 Figs 27-30. Gene Set Level Functional Coherence - CLEAN Scores**.

**S1 Figs 31-38. Gene Set Level Functional Coherence & Enriched Gene Sets - Heatmaps**.

**S1 Figs 39-41. Gene Set Level Benchmark Dose (BMD) Coherence - Heatmaps**.

**S1 Tables 6-13. Core Gene Comparison Tables**.

