## Supplementary material for "ALOHA: Aggregated local extrema splines for high-throughput dose-response analysis": S1 Supplemental Material

#### 1 Available Genomic Dose-Response Methods

| Modeling Features | ALOHA | BMDExpress [1–3] | tcplFit [4, 5] | Reynolds et al. (2020) [6] | GRAVEE [7] |
|---|---|---|---|---|---|
| Differential Expression Pre-filtering | | X | X | | |
| Non-monotonic Fits | X | | | X | |
| Gene Level Fits | X | X | | X | |
| Gene Level BMDs | X | X | | X | |
| Gene Set Level Trends | X | | X | | X |
| Gene Set Level BMDs | X | X | X | | X |
| Parametric Models | | X | X | | |
| Non-parametric Models | X | | | X | X |
| Bayesian Modeling | X | | | X | |
| Frequentist Modeling | | X | X | | X |
| *A Priori* Gene Groups | | X | X | | X |
| Self-Organizing Gene Groups | X | | | | |

**Table 1.** Comparison of features in various genomic dose-response (GDR) methods. Different GDR modeling methods are listed in the columns and rows list key features in the modeling methods. "X" indicates a given GDR method has the listed feature. (BMD = benchmark dose estimate).

#### 2 Comparison Outline Supplement

The comparison between the the two genomic dose-response (GDR) modeling approaches – BMDExpress and ALOHA – described the paper is summarized by the following figure.

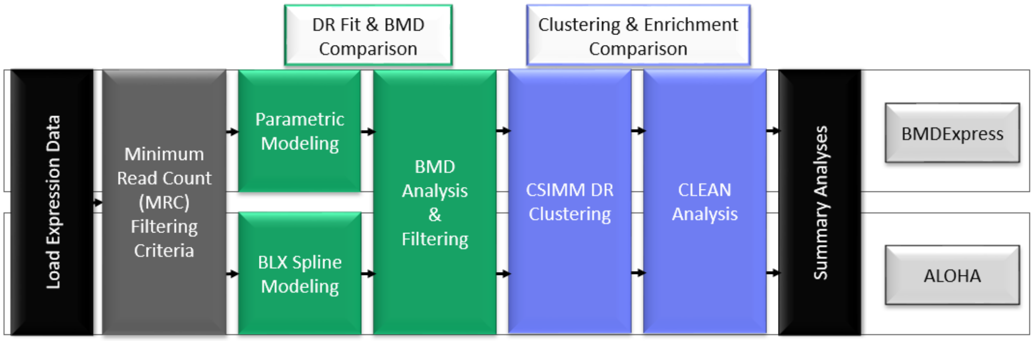


**Fig 1.** Workflow chart outlining our key comparative analyses for evaluating our GDR modeling approach ALOHA in comparison with BMDExpress – adapted from the "Basic Workflow" figure in Auerbach [1].

#### 3 Methods Supplement Material

##### 3.1 BLX Spline Curve Shapes (Extrema = 2 & Increasing = TRUE)

Example BLX spline curve shapes and posterior samples for changepoints. Location of posterior samples of the two change-points in the parameter space with respect to each other provides information about the resulting curve shape from the BLX spline fit. The following equations were used to simulate the five possible curve shapes fit by the BLX spline specifications mentioned in the main paper in §0.1.1:

$$g_1(x) = 2.75x + 6 \tag{1}$$
$$g_2(x) = -2.5x + 15 \tag{2}$$
$$g_3(x) = 7sin(0.3\pi x) + 15 \tag{3}$$
$$g_4(x) = 2.3(x - 1.245)^2 \tag{4}$$
$$g_5(x) = 5.75cos(0.06\pi x - 5) + (x + 5)^1.4 \tag{5}$$
$$g_6(x) = 7.75 + 0.05x + 0.002x^2 \tag{6}$$

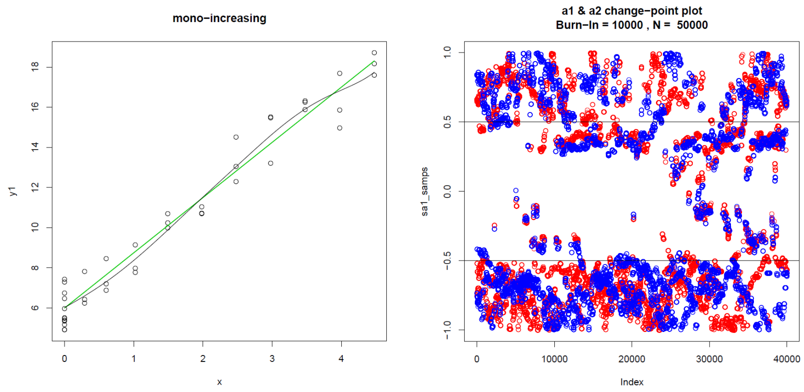


**(a)** **(b)**

**Fig 2.** BLX spline fit (black line) to monotonic increasing trend $g_1(x)$ (green line). (Left) The observed data points and BLX spline estimation. (Right) The MCMC chain of the posterior samples for both change points (i.e. $\alpha_1$ in red and $\alpha_2$ in blue).

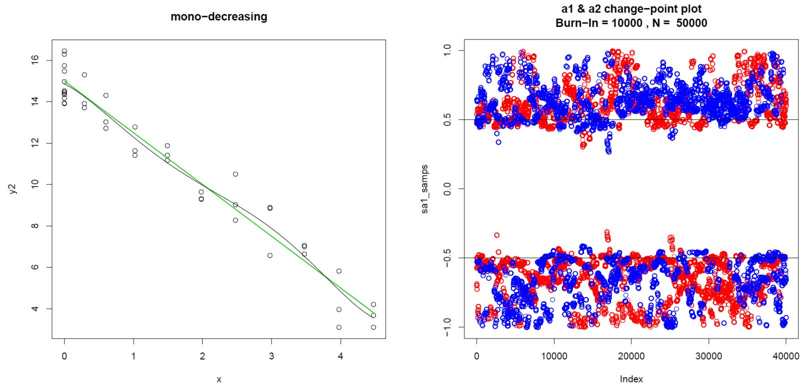


**(a)** **(b)**

**Fig 3.** BLX spline fit (black line) to monotonic decreasing trend $g_2(x)$ (green line). (Left) The observed data points and BLX spline estimation. (Right) The MCMC chain of the posterior samples for both change points (i.e. $\alpha_1$ in red and $\alpha_2$ in blue).

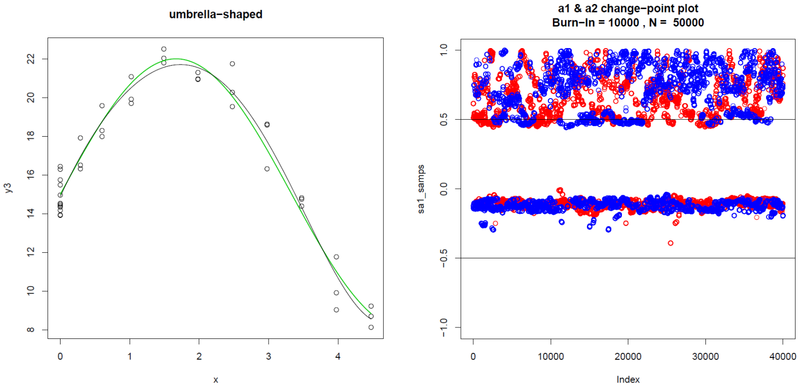


**(a)** **(b)**

**Fig 4.** BLX spline fit (black line) to umbrella-shaped trend $g_3(x)$ (green line). (Left) The observed data points and BLX spline estimation. (Right) The MCMC chain of the posterior samples for both change points (i.e. $\alpha_1$ in red and $\alpha_2$ in blue).

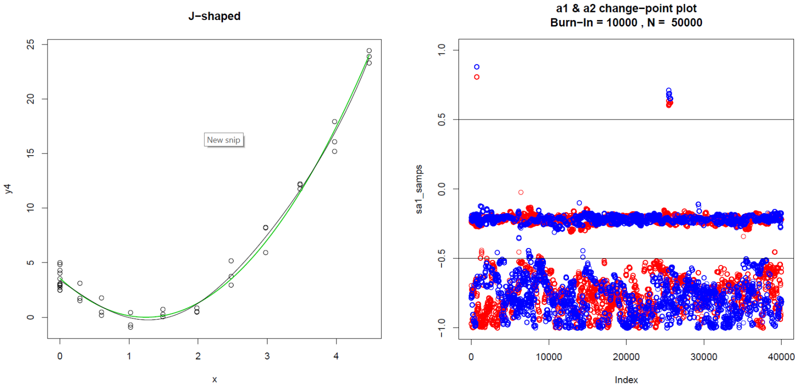


**(a)** **(b)**

**Fig 5.** BLX spline fit (black line) to J-shaped trend $g_4(x)$ (green line). (Left) The observed data points and BLX spline estimation. (Right) The MCMC chain of the posterior samples for both change points (i.e. $\alpha_1$ in red and $\alpha_2$ in blue).

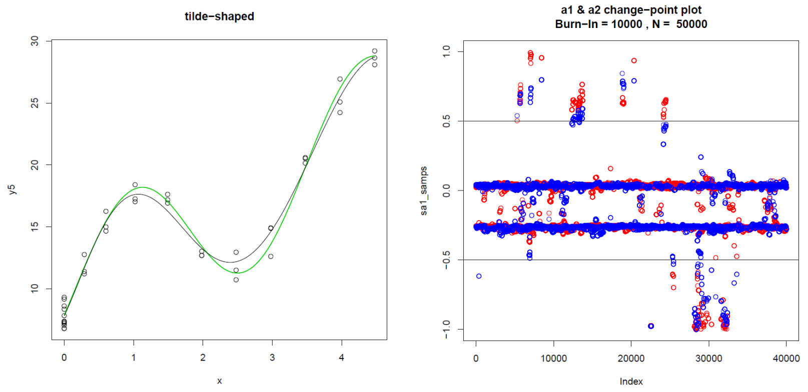


**(a)** **(b)**

**Fig 6.** BLX spline fit (black line) to tilde-shaped trend $g_5(x)$ (green line). (Left) The observed data points and BLX spline estimation. (Right) The MCMC chain of the posterior samples for both change points (i.e. $\alpha_1$ in red and $\alpha_2$ in blue).

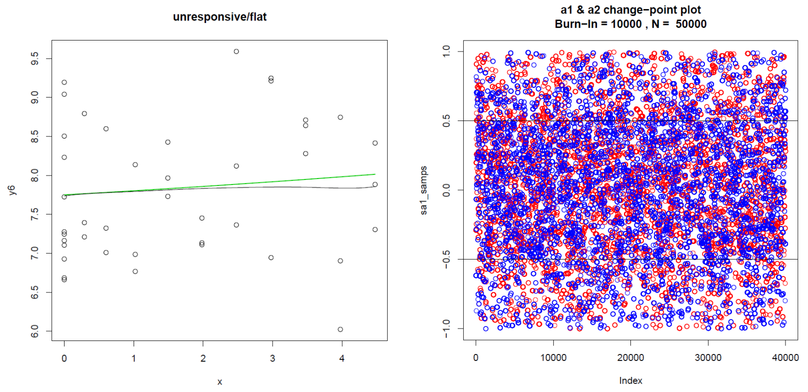


**(a)** **(b)**

**Fig 7.** BLX spline fit (black line) to unresponsive/flat trend $g_6(x)$ (green line). (Left) The observed data points and BLX spline estimation. (Right) The MCMC chain of the posterior samples for both change points (i.e. $\alpha_1$ in red and $\alpha_2$ in blue).

##### 3.2 BMD Estimation BLX Spline Modeling

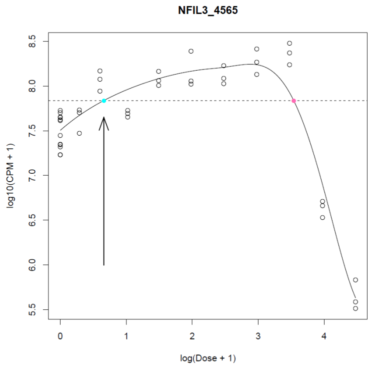


**Fig 8.** Example probe from BAP (AD) with an umbrella-shaped DR curve illustrating BMD estimation for BLX spline estimating umbrella or J-shaped DR curves (i.e. BMR obtains 2 possible BMD estimates). The dotted horizontal line is the BMR value and the aqua blue and pink dots are the potential BMD estimates. The arrow pointing to the aqua blue dot indicates the dose we chose to be the BMD estimate for this curve.

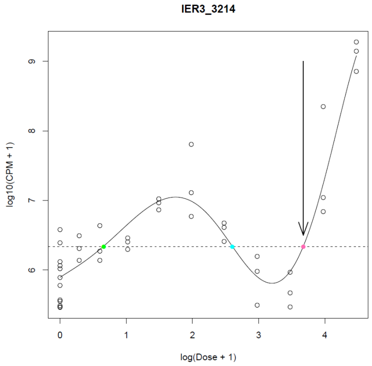


**Fig 9.** Example probe from BAP (AD) with an umbrella-shaped DR curve illustrating BMD estimation for BLX spline estimating tilde-shaped DR curves (i.e. BMR obtains 3 possible BMD estimates). The dotted horizontal line is the BMR value and the green, aqua blue, and pink dots are the potential BMD estimates. The arrow pointing to the pink dot indicates the dose we chose to be the BMD estimate for this curve.

##### 3.3 CSIMM Gene Clustering

Previously, we observed $n$ genes; assume $n^* \leq n$ of these genes can be clustered. That is, after the fitting process $n^*$ genes have acceptable fits. Consider each sample $j \in \{1, \ldots, p\}$ falls into one of $M$ experimental contexts. Here, we specify the contexts as their respective dose groups (i.e. $M = D$). As before, let $Y$ be the $n^* \times p$ matrix of observations we are clustering. Each gene $i$ is evaluated for membership to global and local clusters via a MCMC algorithm.

For each iteration, CSIMM first evaluates the probability $y_i$ is a member of the global cluster $q \in \{1, \ldots, Q\}$ described as the multivariate-normal (MVN):

$$y_i \sim N(\mu_q, \Sigma_q) \tag{7}$$

where $\mu_q$ is the mean vector and $\Sigma_q$ is the variance-covariance matrix for global cluster $q$. Once all genes are assigned to a global cluster, each global cluster $q$ is evaluated for membership to a local cluster in some experimental context $f \in \{1, \ldots, M\}$. Without loss of generality, for context $f$ CSIMM evaluates the probability that $\mu_{(q,f)}$ – a sub-vector of global cluster $q$ – belongs to a local cluster $w$. The local cluster $w$ is described by the MVN:

$$\mu_{(q,f)} \sim N(\mu_w^*, \Sigma_w^*) \tag{8}$$

with the mean vector $\mu_w^*$ and the context-specific variance-covariance matrix $\Sigma_w^*$.

After genes are assigned to global and local clusters, parameters for all global MVNs, all local MVNs, and the hyperparameters are updated based on membership in each cluster.

At the end of the MCMC chain the posterior pairwise probabilities (PPP) is calculated for genes belonging to the same global and local clusters. For example, consider gene $a$ and $b$ are assigned the same group labels for 85% of the MCMC iterations after burn-in, then the PPP for these genes is 0.85. Calculating the PPP of genes avoids finding the "correct" number of resulting gene sets and circumvents the label-switching problem, common with DPMs (see Celeux et al. [8] and Stephens [9], among others). PPP's are then used to construct a hierarchical tree using 1-PPP as the distance measure. Using 1-PPP as the distance measure this places gene sets with a high PPP closer to the leaves of the tree and are obtained with smaller cut-off values (e.g. if PPP = 0.95, then h = 0.05).

### 4 Results Supplement Material

#### 4.1 Fit Comparison for Cases where Parametric Fit Passes but BLX Spline Fails to Pass

**Case 1: Noisy expression Response Data.**

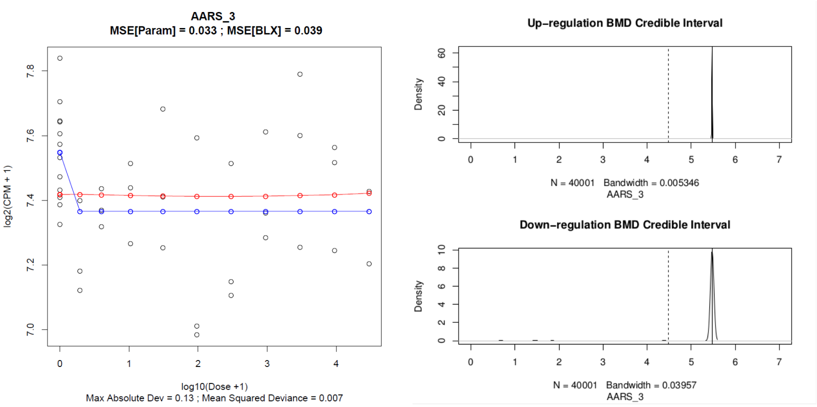


**(a)** **(b)**

**Fig 10.** (Left) Plot containing the best parametric (blue) and BLX spline (red) fits on the observed expression response (points) for the "AARS_3" probe from the BAP-AD dataset. (Right) The BLX spline BMD posterior distribution plots, for both the up (top) and down (bottom) regulation directions. The black dotted line indicates the maximum experimental dose on the $log_{10}(Dose + 1)$ scale. Both regulation directions have a BMD posterior distribution highly concentrated around 1 plus the maximum experimental dose, as indicated by the sharp peaks right of the black dotted line. The credible interval, for both regulations directions, is indicated by a solid black like located at 1 plus the maximum experimental dose ($BMD_{95\%CI} = (5.48, 5.48)$). The down-regulation BMD estimate plot (bottom-right) provides the best illustration.

**Case 2: Active and Inactive Samples in the Cytotoxic Dose Groups.**

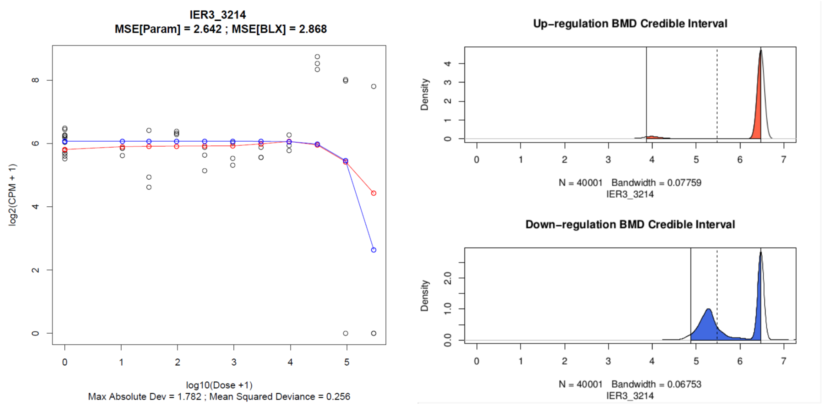


(a) (b)

**Fig 11.** (Left) Plot containing the best parametric (blue) and BLX spline (red) fits on the observed expression response (points) for the "IER3_3214" probe from the CPZ-AD dataset. (Right) The BLX spline BMD posterior distribution plots, for both the up (top) and down (bottom) regulation directions. The black dotted line indicates the maximum experimental dose on the $log_{10}(Dose + 1)$ scale. Credible intervals are indicated by the red (up-regulation – $BMD_{95\%CI} = (3.87, 6.48)$) and blue (down-regulation – $BMD_{95\%CI} = (4.88, 6.48)$) shaded regions.

**Case 3: Unreliable (or Wide) BLX spline BMD credible intervals Otherwise Seemingly Active.**

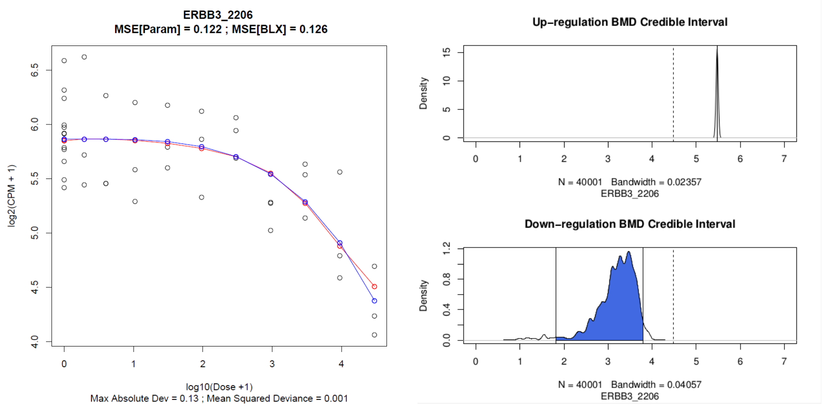


(a) (b)

**Fig 12.** (Left) Plot containing the best parametric (blue) and BLX spline (red) fits on the observed expression response (points) for the "ERBB3_2206" probe from the BAP-AD dataset. (Right) The BLX spline BMD posterior distribution plots, for both the up (top) and down (bottom) regulation directions. The black dotted line indicates the maximum experimental dose on the $log_{10}(Dose + 1)$ scale. In the top-right plot, the credible interval for the BMD estimate in the up-regulation direction is indicated by a solid black line at 1 plus the maximum dose ($BMD_{95\%CI} = (5.48, 5.48)$). In the bottom-right plot, the blue shaded region indicates the credible interval for the BMD estimate in the down-regulation direction ($BMD_{95\%CI} = (1.81, 3.79)$).

**Case 4: Bi-modal BLX spline BMD credible intervals with at Least One**

**Mode Greater than the Maximum Dose.**

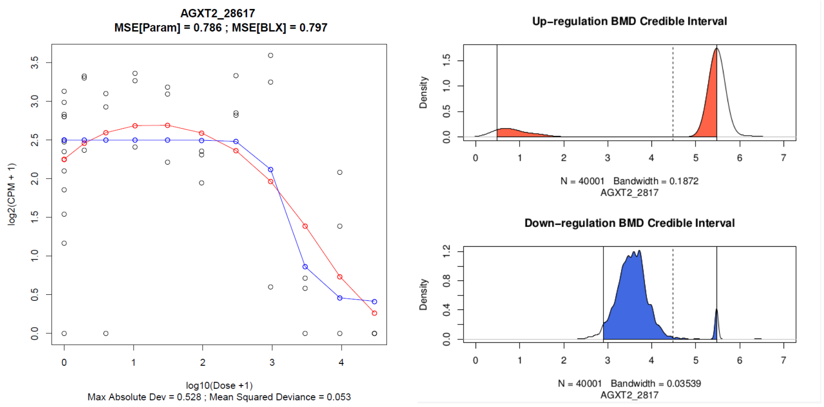


**(a)** **(b)**

**Fig 13.** (Left) Plot containing the best parametric (blue) and BLX spline (red) fits on the observed expression response (points) for the "AGXT2_28617" probe from the BAP-AD dataset. (Right) The BLX spline BMD posterior distribution plots, for both the up (top) and down (bottom) regulation directions. The black dotted line indicates the maximum experimental dose on the $log_{10}(Dose + 1)$ scale. Credible intervals are indicated by the red (up-regulation – $BMD_{95\%CI} = (0.48, 5.48)$) and blue (down-regulation – $BMD_{95\%CI} = (2.89, 5.48)$) shaded regions.

#### 4.2 BMD Estimate Comparison

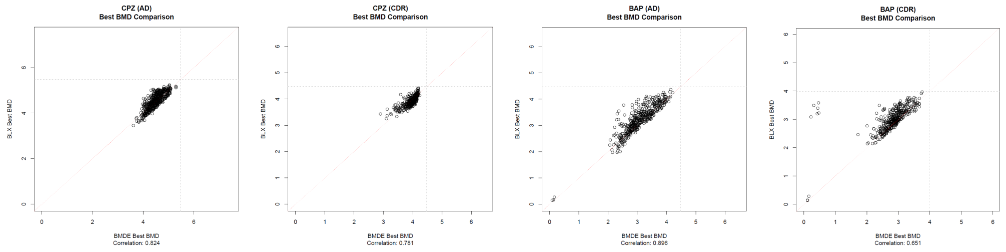


**(a)** **(b)** **(c)** **(d)**

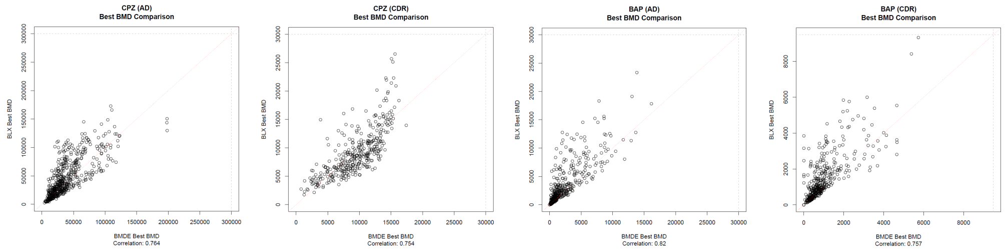


**(e)** **(f)** **(g)** **(h)**

**Fig 14.** Scatterplot of the best BMD estimates from the best parametric fit (x-axis) and the BLX spline fit (y-axis) for all datasets, only for those probes with the best BMD estimate less than the maximum experimental dose and BMDU/BMDL less than 40 in both BMDExpress (version 2.3) and the BLX spline modeling. (Top row) Logarithmic scale BMD values. (Bottom row) Original dose scale BMD values in $nM$.

##### Chlorpromazine All Doses

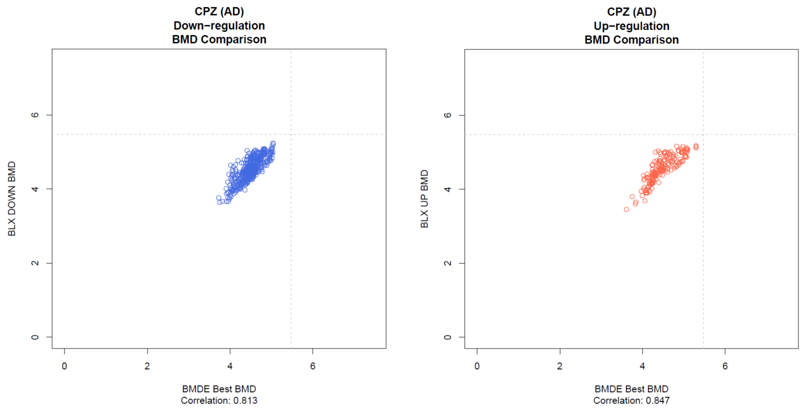


**(a)** Down-regulation **(b)** Up-regulation

**Fig 15.** Scatterplot of the BMD estimates from the best parametric fit (x-axis) and the BLX spline fit (y-axis), based on the regulation direction from BMDExpress (version 2.3), for CPZ all doses, only for those probes passing "reliability" criteria in both BMDExpress and the BLX spline modeling. (Left) Down-regulated BMD estimate comparison, only probes where BMDExpress adverse direction = -1 (down-regulation). (Right) Up-regulation BMD estimate comparison, only probes where BMDExpress adverse direction = 1 (up-regulation).

##### Chlorpromazine Cytotoxic Doses Removed

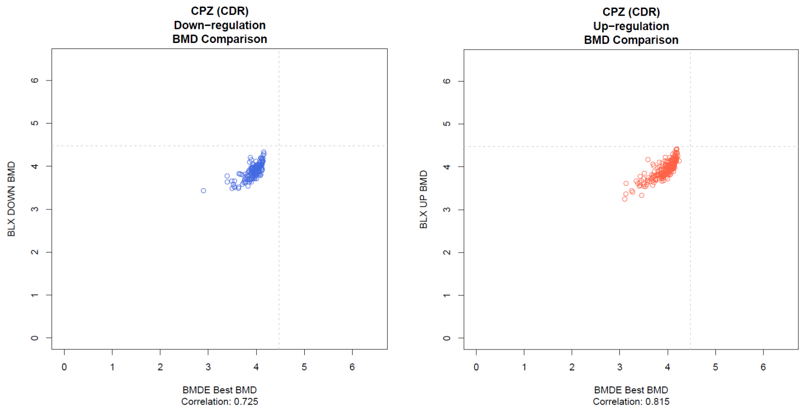


**(a)** Down-regulation **(b)** Up-regulation

**Fig 16.** Scatterplot of the BMD estimates from the best parametric fit (x-axis) and the BLX spline fit (y-axis), based on the regulation direction from BMDExpress (version 2.3), for CPZ cytotoxic doses removed, only for those probes passing "reliability" criteria in both BMDExpress and the BLX spline modeling. (Left) Down-regulated BMD estimate comparison, only probes where BMDExpress adverse direction = -1 (down-regulation). (Right) Up-regulation BMD estimate comparison, only probes where BMDExpress adverse direction = 1 (up-regulation).

##### Benzo-[$\alpha$]-pyrene All Doses

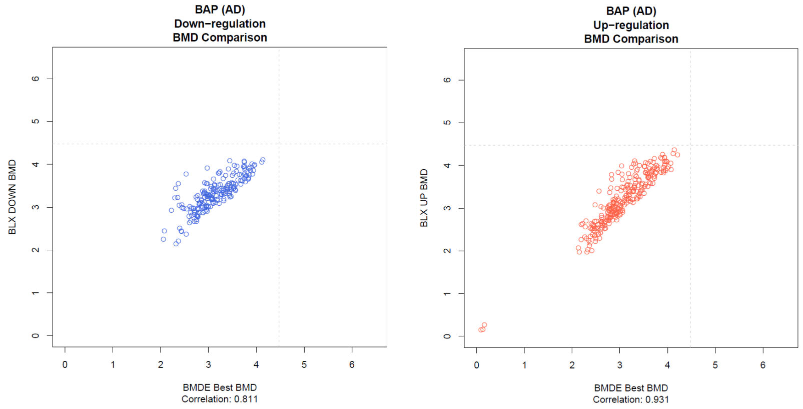


**(a)** Down-regulation **(b)** Up-regulation

**Fig 17.** Scatterplot of the BMD estimates from the best parametric fit (x-axis) and the BLX spline fit (y-axis), based on the regulation direction from BMDExpress (version 2.3), for BAP all doses, only for those probes passing "reliability" criteria in both BMDExpress and the BLX spline modeling. (Left) Down-regulated BMD estimate comparison, only probes where BMDExpress adverse direction = -1 (down-regulation). (Right) Up-regulation BMD estimate comparison, only probes where BMDExpress adverse direction = 1 (up-regulation).

###### Benzo-[$\alpha$]-pyrene Cytotoxic Doses Removed

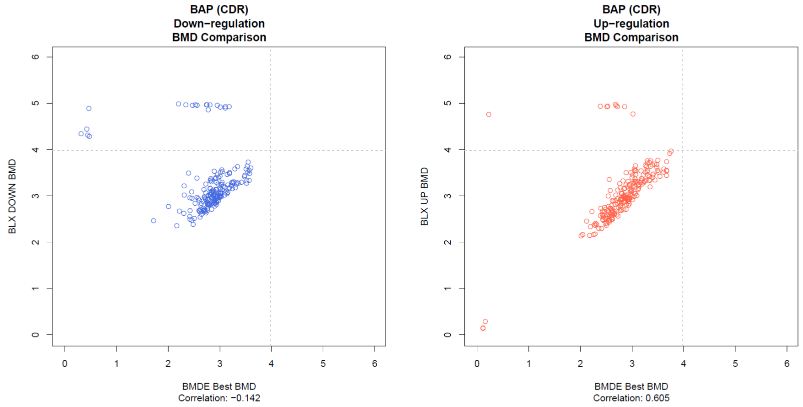


**(a)** Down-regulation **(b)** Up-regulation

**Fig 18.** Scatterplot of the BMD estimates from the best parametric fit (x-axis) and the BLX spline fit (y-axis), based on the regulation direction from BMDExpress (version 2.3), for BAP cytotoxic doses removed, only for those probes passing "reliability" criteria in both BMDExpress and the BLX spline modeling. (Left) Down-regulated BMD estimate comparison, only probes where BMDExpress adverse direction = -1 (down-regulation). (Right) Up-regulation BMD estimate comparison, only probes where BMDExpress adverse direction = 1 (up-regulation).

#### 4.3 Dose-Response Clustering Comparison

**Plot description:** Heatmaps of the CSIMM clusters (left column) and ALOHA clusters (right column) for all datasets assessed. (From left to right in each heatmap plot) The main plot displays the normalized expression values with all replicates. Next are the heatmaps displaying BLX spline parameter estimates, i.e. change-points and spline coefficients respectively. Finally, down and up regulated mean BMD estimates for each probe are displayed in the right-most heatmap (BMDs above the maximum dose are NA's and are displayed in black).

**Chlorpromazine All Doses**

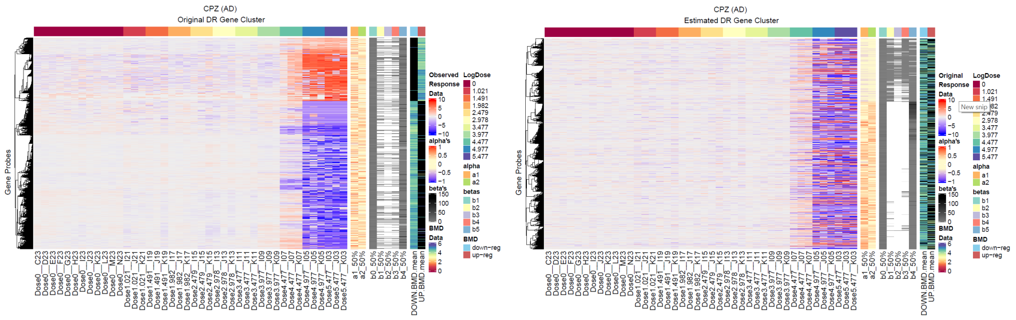


**(a)** CSIMM **(b)** ALOHA

**Fig 19. CPZ AD** - Heatmaps displaying clustering results applied to normalized expression data showing visual separation of DR trends. (Left - *CSIMM*) CSIMM clustering on normalized expression data with all replicates in each dose group. (Right - *ALOHA*) CSIMM clustering on estimated response for each experimental dose group.

**Chlorpromazine Cytotoxic Doses Removed**

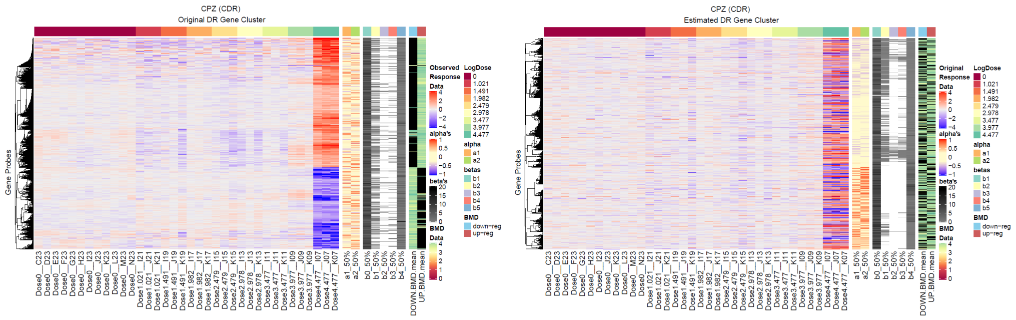


**(a)** CSIMM **(b)** ALOHA

**Fig 20. CPZ CDR** - Heatmaps displaying clustering results applied to normalized expression data showing visual separation of DR trends. (Left - *CSIMM*) CSIMM clustering on normalized expression data with all replicates in each dose group. (Right - *ALOHA*) CSIMM clustering on estimated response for each experimental dose group.

**Benzo-[$\alpha$]-pyrene All Doses**

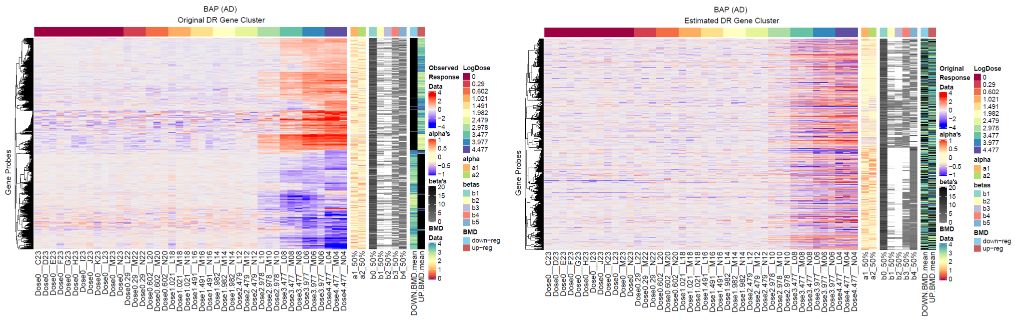


**(a)** CSIMM **(b)** ALOHA

**Fig 21. BAP AD** - Heatmaps displaying clustering results applied to normalized expression data showing visual separation of DR trends. (Left - *CSIMM*) CSIMM clustering on normalized expression data with all replicates in each dose group. (Right - *ALOHA*) CSIMM clustering on estimated response for each experimental dose group.

**Benzo-[$\alpha$]-pyrene Cytotoxic Doses Removed**

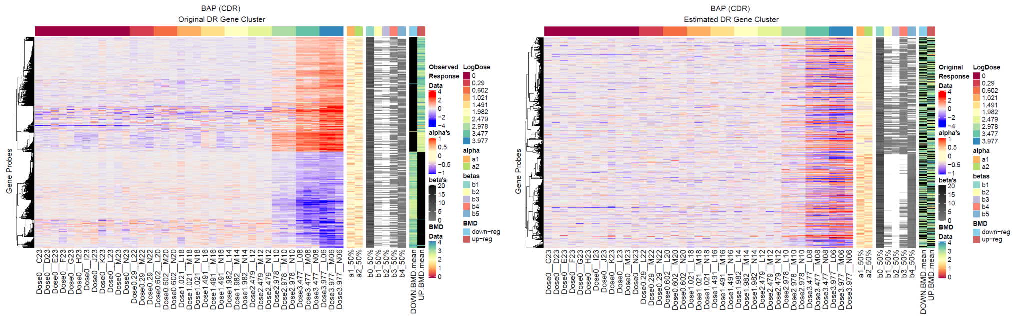


**(a)** CSIMM **(b)** ALOHA

**Fig 22. BAP CDR** - Heatmaps displaying clustering results applied to normalized expression data showing visual separation of DR trends. (Left - *CSIMM*) CSIMM clustering on normalized expression data with all replicates in each dose group. (Right - *ALOHA*) CSIMM clustering on estimated response for each experimental dose group.

**Plot description:** Heatmaps of the CSIMM clusters (left column) and ALOHA clusters (right column) for all datasets assessed. (From left to right in each heatmap plot) The main plot displays the BLX estimated dose-response data. The posterior pairwise probability dendrogram is cut at the height resulting in five clusters. The chosen cut-off was arbitrary, since there many possible cut-offs exist when evaluating clustering approaches that result in a dendrogram, but this was done for exposition of dose-response coherence within resulting clusters from the CSIMM and ALOHA algorithms.

##### Chlorpromazine All Doses

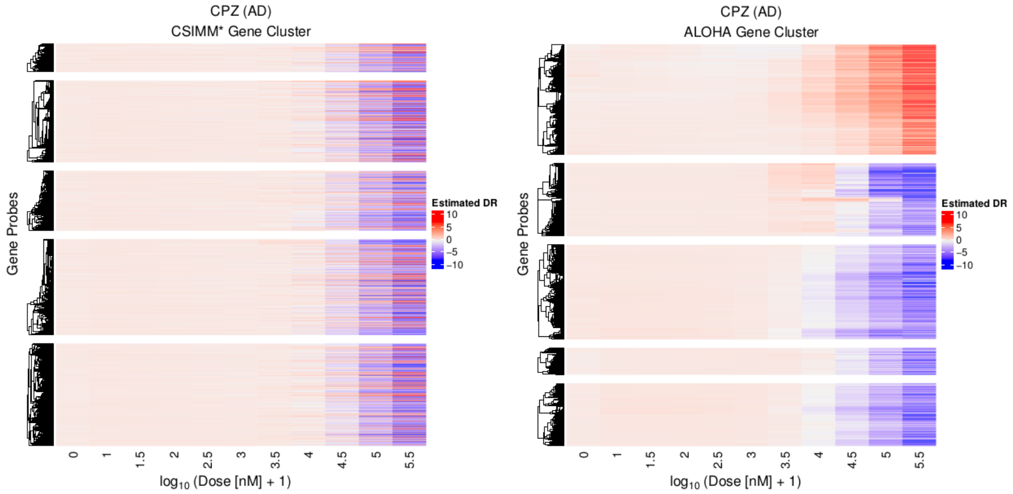


**(a)** CSIMM **(b)** ALOHA

**Fig 23. CPZ AD** - Heatmaps displaying clustering results applied to BLX estimated dose-response data showing visual separation of DR trends. (Left - *CSIMM*) CSIMM clustering on observed normalized expression data with all replicates in each dose group. (Right - *ALOHA*) CSIMM clustering on BLX estimated dose-response data.

| Chlorpromazine (AD) | ALOHA | CSIMM |
|---|---|---|
| Avg. Distance Between Clusters | 8.66 | 7.35 |
| Avg. Distance Within Clusters | 2.97 | 7.33 |
| Dunn Index | 6.0e-3 | 9.5e-4 |
| Avg. Silhouette Width | 0.19 | -0.07 |
| Entropy | 1.52 | 1.53 |

**Table 2.** Scores measuring the internal coherence of dose-response trends in the five clusters resulting from ALOHA and CSIMM in the Chlorpromazine dataset with all doses, see Fig 23. Larger values for average distance between clusters, Dunn index, and the average silhouette width indicate better separation of genes into similar dose-response groups. Smaller values for the average distance within clusters and entropy indicate more coherence between genes clustered together.

#### Chlorpromazine Cytotoxic Doses Removed

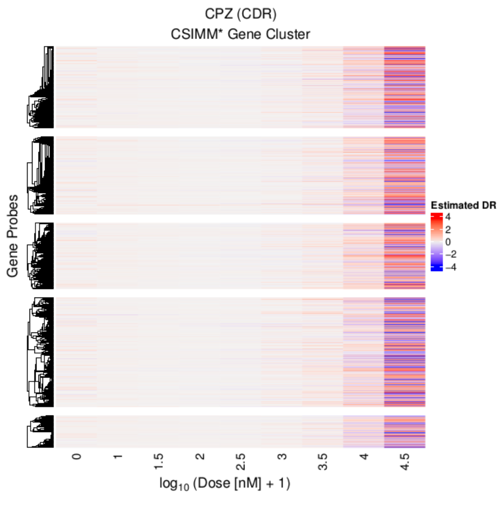


(a) CSIMM

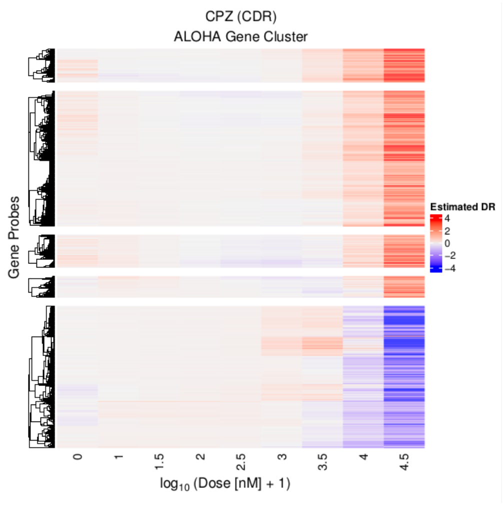


(b) ALOHA

**Fig 24. CPZ CDR** - Heatmaps displaying clustering results applied to BLX estimated dose-response data showing visual separation of DR trends. (Left - *CSIMM*) CSIMM clustering on observed normalized expression data with all replicates in each dose group. (Right - *ALOHA*) CSIMM clustering on BLX estimated dose-response data.

| Chlorpromazine (CDR) | ALOHA | CSIMM |
|---|---|---|
| Avg. Distance Between Clusters | 3.85 | 3.11 |
| Avg. Distance Within Clusters | 1.42 | 3.10 |
| Dunn Index | 3.03-3 | 1.1e-3 |
| Avg. Silhouette Width | 0.23 | -0.10 |
| Entropy | 1.34 | 1.55 |

**Table 3.** Scores measuring the internal coherence of dose-response trends in the five clusters resulting from ALOHA and CSIMM in the Chlorpromazine dataset without cytotoxic doses, see Supplementary Fig 24. Larger values for average distance between clusters, Dunn index, and the average silhouette width indicate better separation of genes into similar dose-response groups. Smaller values for the average distance within clusters and entropy indicate more coherence between genes clustered together.

##### Benzo-[$\alpha$]-pyrene All Doses

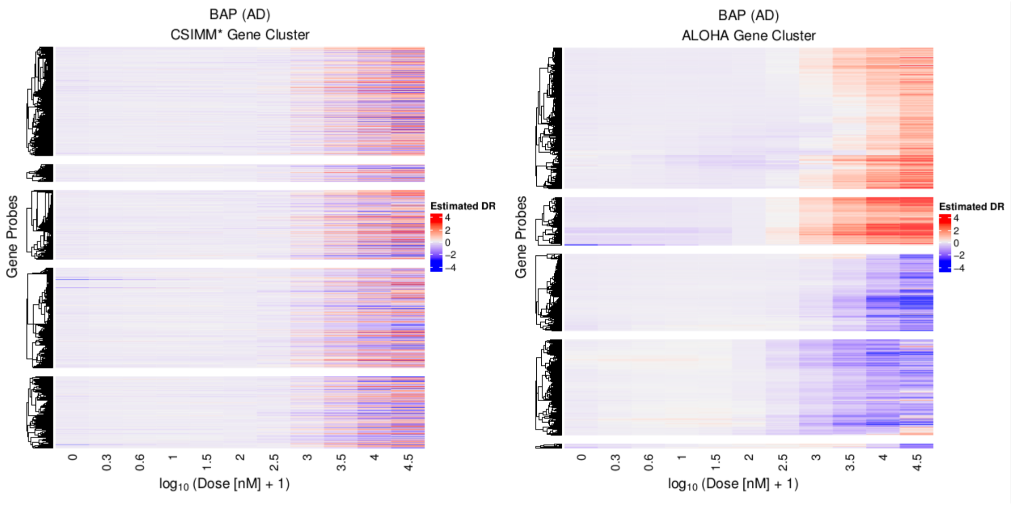


**(a)** CSIMM **(b)** ALOHA

**Fig 25. BAP AD** - Heatmaps displaying clustering results applied to BLX estimated dose-response data showing visual separation of DR trends. (Left - *CSIMM*) CSIMM clustering on observed normalized expression data with all replicates in each dose group. (Right - *ALOHA*) CSIMM clustering on BLX estimated dose-response data.

| Benzo-[$\alpha$]-pyrene (AD) | ALOHA | CSIMM |
|---|---|---|
| Avg. Distance Between Clusters | 3.84 | 3.23 |
| Avg. Distance Within Clusters | 1.68 | 3.21 |
| Dunn Index | 4.4e-3 | 9.3e-4 |
| Avg. Silhouette Width | 0.20 | -0.06 |
| Entropy | 1.37 | 1.49 |

**Table 4.** Scores measuring the internal coherence of dose-response trends in the five clusters resulting from ALOHA and CSIMM in the Benzo-[$\alpha$]-pyrene dataset with all doses, see Fig 25. Larger values for average distance between clusters, Dunn index, and the average silhouette width indicate better separation of genes into similar dose-response groups. Smaller values for the average distance within clusters and entropy indicate more coherence between genes clustered together.

**Benzo-[$\alpha$]-pyrene Cytotoxic Doses Removed**

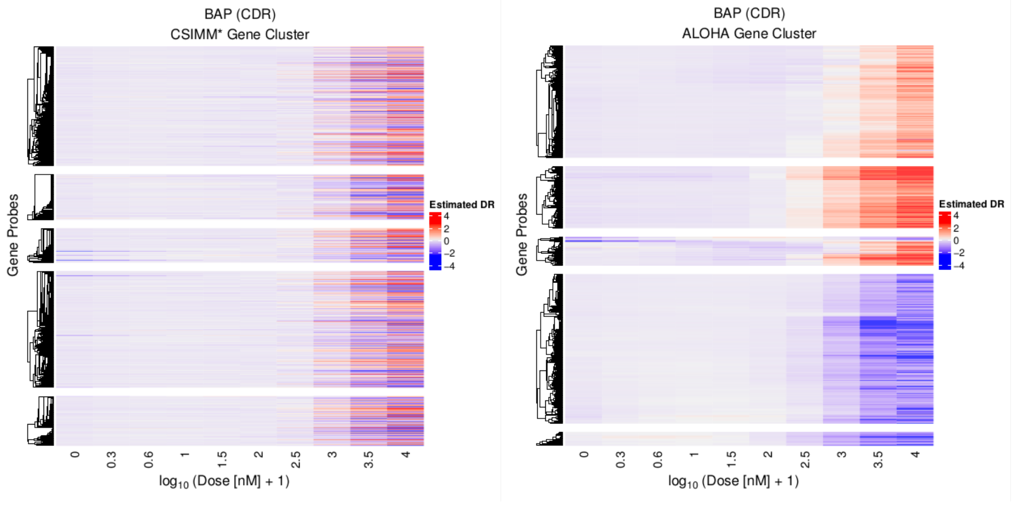


**(a)** CSIMM **(b)** ALOHA

**Fig 26. BAP CDR** - Heatmaps displaying clustering results applied to BLX estimated dose-response data showing visual separation of DR trends. (Left - *CSIMM*) CSIMM clustering on observed normalized expression data with all replicates in each dose group. (Right - *ALOHA*) CSIMM clustering on BLX estimated dose-response data.

| Benzo-[$\alpha$]-pyrene (CDR) | ALOHA | CSIMM |
|---|---|---|
| Avg. Distance Between Clusters | 3.21 | 2.59 |
| Avg. Distance Within Clusters | 1.11 | 2.57 |
| Dunn Index | 5.6e-3 | 1.5e-3 |
| Avg. Silhouette Width | 0.23 | -0.04 |
| Entropy | 1.34 | 1.48 |

**Table 5.** Scores measuring the internal coherence of dose-response trends in the five clusters resulting from ALOHA and CSIMM in the Benzo-[$\alpha$]-pyrene dataset without cytotoxic doses, see Supplementary Fig 26. Larger values for average distance between clusters, Dunn index, and the average silhouette width indicate better separation of genes into similar dose-response groups. Smaller values for the average distance within clusters and entropy indicate more coherence between genes clustered together.

#### 4.4 Gene Level Functional Coherence

Figs 27-30 compare the empirical cumulative distribution function (ECDF) of CLEAN scores for each strategy on all the datasets. Functional coherence is indicated by ECDF plots with high CLEAN scores (i.e. slowest to reach 1 and furthest to the right on the ECDF plot). In general, the BMDExpress approach (blue) showed the lowest level of functional coherence. For the other two approaches, other than Chlorpromazine with all doses, there is little separation between the ECDF curves. For Chlorpromazine without cytotoxic doses, there was greater separation at the highest scores; however, these constitute less than 5% of the genes, and subsequently, it may not indicate a difference between the approaches. For Chlorpromazine with all doses, the ECDF curves for the Hallmark gene set overlap indicating the methods are equivocal. Using the Cancer Gene Neighborhood gene list, there were clear improvements using the CSIMM approach.

Overall, similar levels of functional coherence were seen using CSIMM and ALOHA. However, since the genes are grouped differently, as described in §3.2.1 incorporation of the model fit data does not reduce our ability to obtain biologically relevant clusters. The flexibility of the BLX spline obtains more accurate fits and when coupled with clustering helps identify biologically relevant clusters.

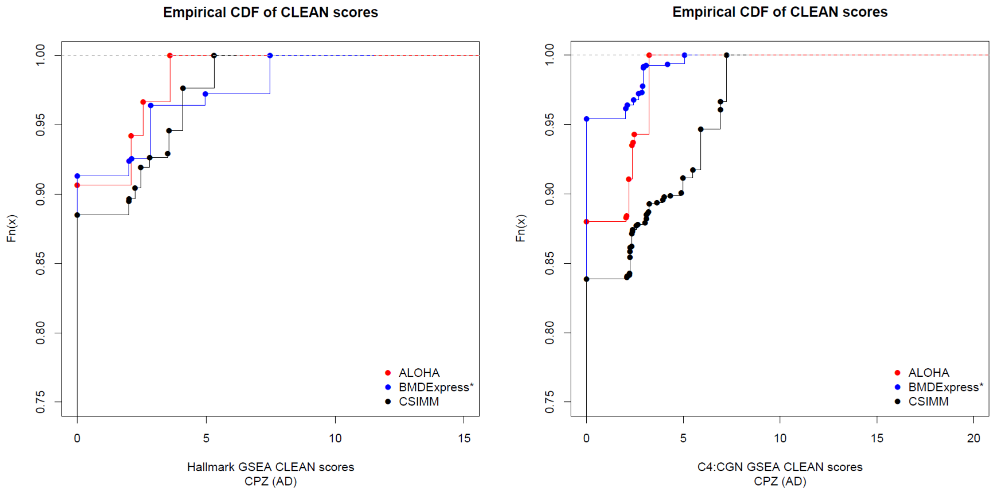


**(a)** Hallmark **(b)** Cancer Gene Neighborhoods

**Fig 27.** CLEAN score empirical cumulative distribution function (ECDF) plots for Chlorpromazine including all doses (CPZ-AD), with MSigDB's Hallmark (right) and Cancer Gene Neighborhood (CGN) (left) gene lists on the normalized expression (CSIMM – black), BLX estimated expression (ALOHA – red), and best parametric fit estimated expression (BMDExpress – blue) approaches.

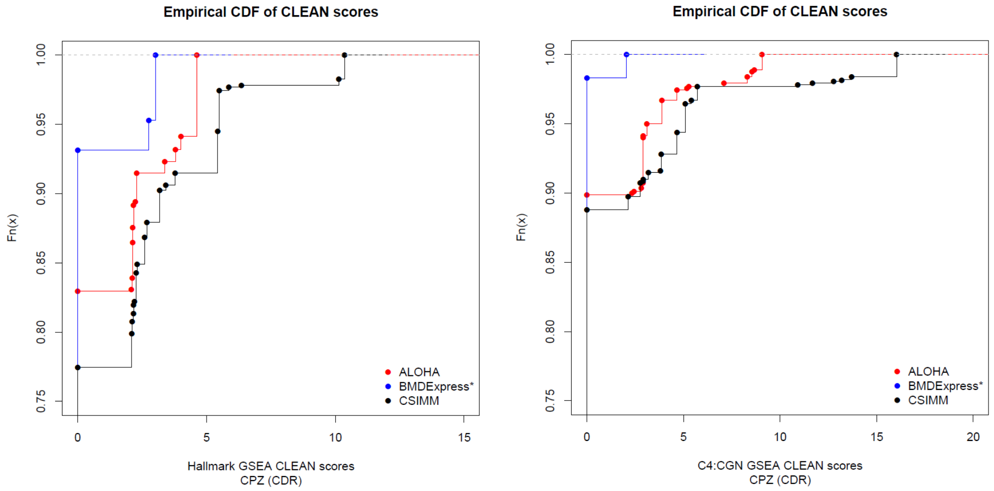


**(a)** Hallmark **(b)** Cancer Gene Neighborhoods

**Fig 28.** CLEAN score empirical cumulative distribution function (ECDF) plots for Chlorpromazine without cytotoxic doses (CPZ-CDR), with MSigDB's Hallmark (right) and Cancer Gene Neighborhood (CGN) (left) gene lists on the normalized expression (CSIMM – black), BLX estimated expression (ALOHA – red), and best parametric fit estimated expression (BMDExpress – blue) approaches.

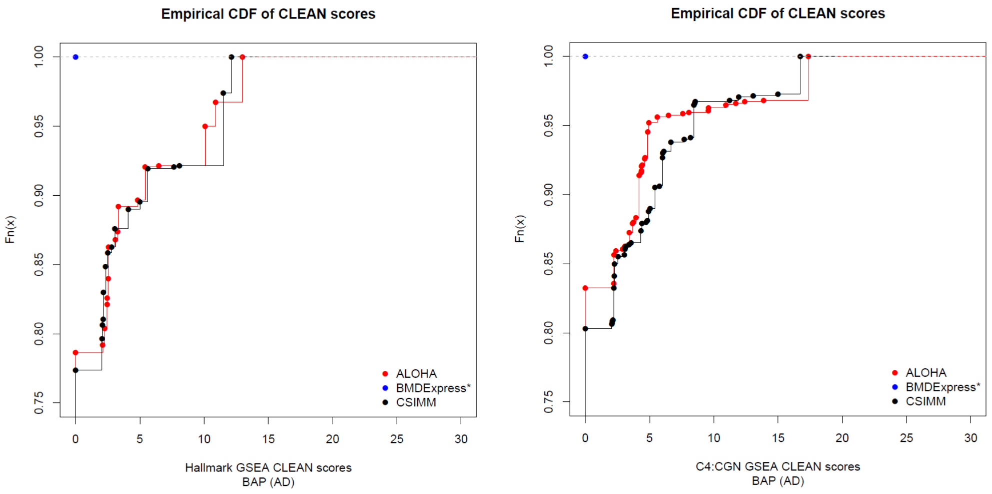


**(a)** Hallmark **(b)** Cancer Gene Neighborhoods

**Fig 29.** CLEAN score empirical cumulative distribution function (ECDF) plots for Benzo-[$\alpha$]-pyrene including all doses (BAP-AD), with MSigDB's Hallmark (right) and Cancer Gene Neighborhood (CGN) (left) gene lists on the normalized expression (CSIMM – black), BLX estimated expression (ALOHA – red), and best parametric fit estimated expression (BMDExpress – blue) approaches.

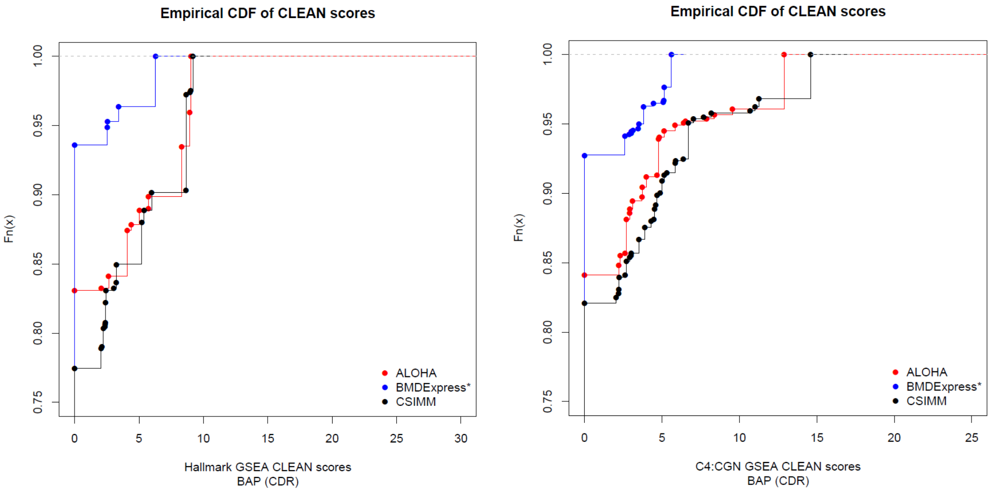


**(a)** Hallmark **(b)** Cancer Gene Neighborhoods

**Fig 30.** CLEAN score empirical cumulative distribution function (ECDF) plots for Benzo-[$\alpha$]-pyrene without cytotoxic doses (BAP-CDR), with MSigDB's Hallmark (right) and Cancer Gene Neighborhood (CGN) (left) gene lists on the normalized expression (CSIMM – black), BLX estimated expression (ALOHA – red), and best parametric fit estimated expression (BMDExpress – blue) approaches.

#### 4.5 Gene Set Level Functional Coherence & Enriched Sets

##### 4.5.1 MSigDB Hallmark Gene Set Significant Categories

**Plot description:** Heatmaps comparing the enrichment scores of the clusters with smallest FDR adjusted Fisher's p-value for the MSigDB Hallmark gene sets found to be significant (FDR adjusted Fisher's p-value $< 0.01$) with the three different data inputs for CSIMM clustering, namely the normalized expression data with all replicates (CSIMM), parametric model estimated response data (BMDExpress), and the BLX spline fit estimated response data (ALOHA). Clusters with the smallest enrichment score not having a minimum of 3 genes mapping to the enriched gene set or populating the gene set by a minimum of 5% were biologically irrelevant due to lack of appropriate information and were removed therefore not displayed in the heatmap. (Left) Heatmap shows $-log_{10}(FisherFDR)$ of each enriched gene set from each approach. (Right) Heatmap shows $log(OddsRatio)$ of each enriched gene set from each approach. Rows are unique Hallmark gene sets and columns indicate clustering approaches.

##### Chlorpromazine All Doses

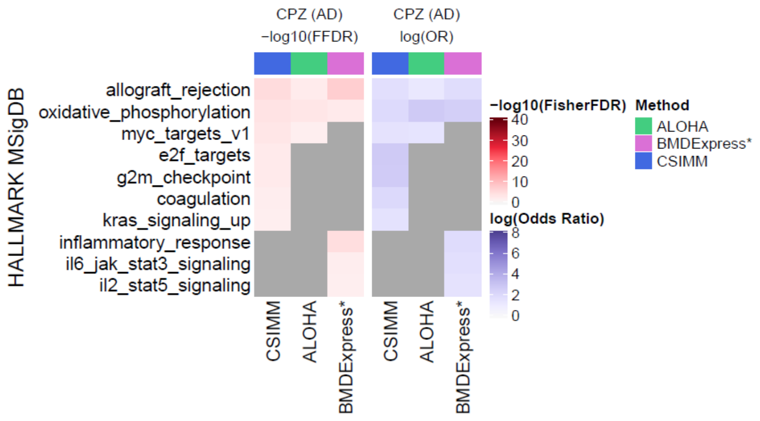


**Fig 31. CPZ AD** Heatmaps of the significant Hallmark gene sets comparing the most statistically significant clusters, displaying negative log-scale Fisher FDR and log odds ratio, respectively.

##### Chlorpromazine Cytotoxic Doses Removed

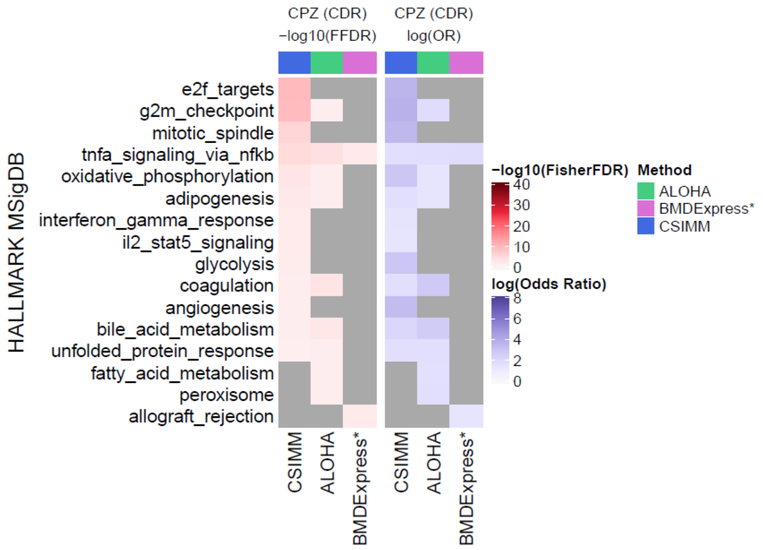


**Fig 32. CPZ CDR** Heatmaps of the significant Hallmark gene sets comparing the most statistically significant clusters, displaying negative log-scale Fisher FDR and log odds ratio, respectively.

##### Benzo-[$\alpha$]-pyrene All Doses

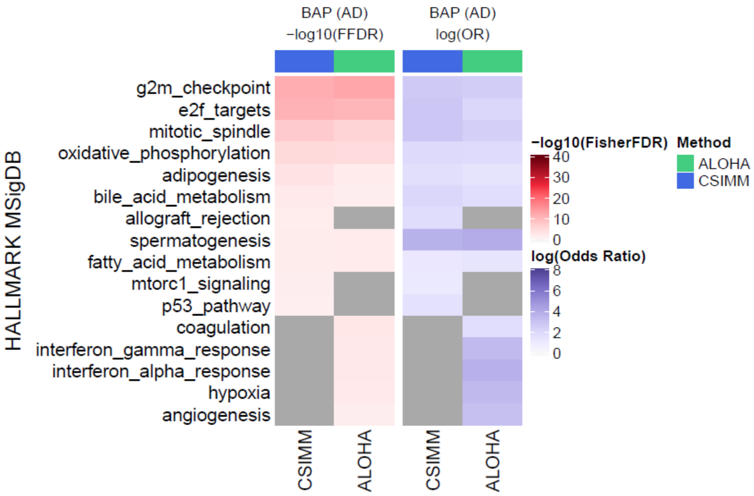


**Fig 33. BAP AD** Heatmaps of the significant Hallmark gene sets comparing the most statistically significant clusters, displaying negative log-scale Fisher FDR and log odds ratio, respectively.

##### Benzo-[$\alpha$]-pyrene Cytotoxic Doses Removed

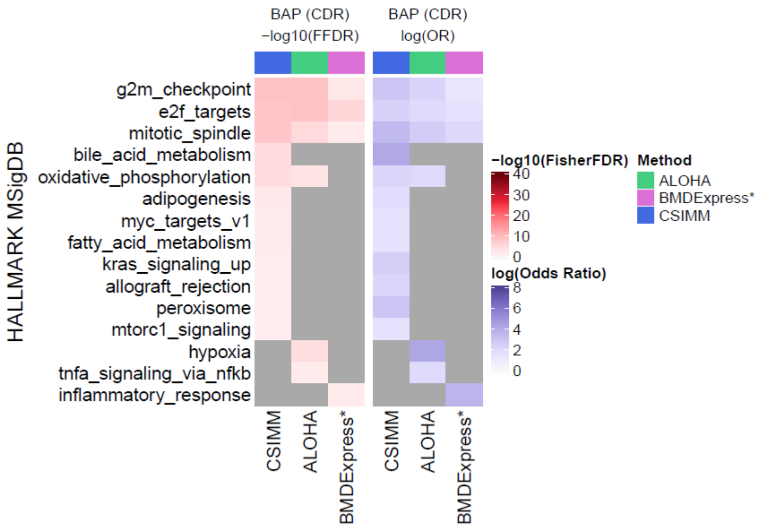


**Fig 34. BAP CDR** Heatmaps of the significant Hallmark gene sets comparing the most statistically significant clusters, displaying negative log-scale Fisher FDR and log odds ratio, respectively.

#### 4.6 MSigDB C4:CGN Gene Set Significant Categories

**Plot description:** Heatmaps comparing the enrichment scores of the clusters with smallest FDR adjusted Fisher's p-value for the MSigDB Computational Cancer Gene Neighborhood gene sets (C4:CGN) found to be significant (FDR adjusted Fisher's p-value $< 0.01$) with the three data inputs, namely the normalized expression data with all replicates (CSIMM), the parametric model estimated response data (BMDExpress*), and the BLX spline fit estimated response data (ALOHA). Clusters with the smallest enrichment score not having a minimum of 3 genes mapping to the enriched gene set or populating the gene set by a minimum of 5% were biologically irrelevant due to lack of appropriate information and removed and are not displayed in the heatmap. (Left) Heatmap shows $-log_{10}(FisherFDR)$ of each enriched gene set from each approach. (Right) Heatmap shows $log(OddsRatio)$ of each enriched gene set from each approach. Rows are unique C4:CGN gene sets and columns indicate clustering approaches.

**Chlorpromazine All Doses**

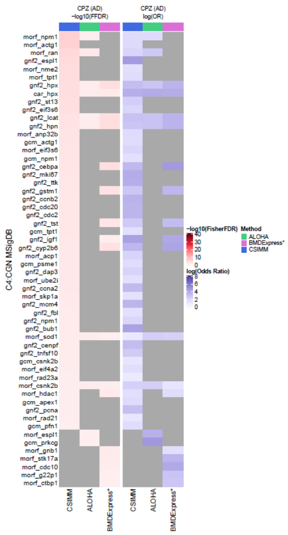


**Fig 35. CPZ AD** Heatmaps of the significant Cancer Gene Neighborhood gene sets comparing the most statistically significant clusters, displaying negative log-scale Fisher FDR and log odds ratio, respectively. Gray indicates which gene sets were not enriched for a given clustering approach.

##### Chlorpromazine Cytotoxic Doses Removed

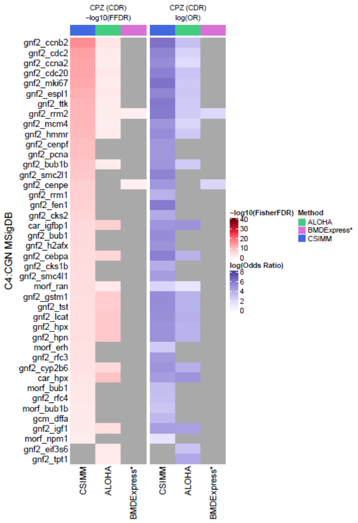


**Fig 36. CPZ CDR** Heatmaps of the significant Cancer Gene Neighborhood gene sets comparing the most statistically significant clusters, displaying negative log-scale Fisher FDR and log odds ratio, respectively. Gray indicates which gene sets were not enriched for a given clustering approach.

##### Benzo-[$\alpha$]-pyrene All Doses

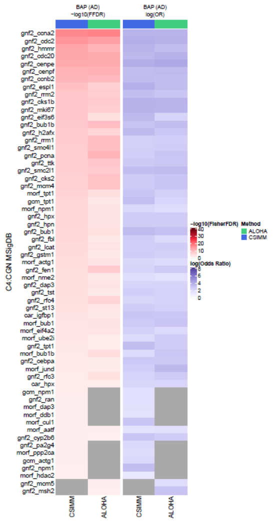


**Fig 37. BAP AD** Heatmaps of the significant Cancer Gene Neighborhood gene sets comparing the most statistically significant clusters, displaying negative log-scale Fisher FDR and log odds ratio, respectively. Gray indicates which gene sets were not enriched for a given clustering approach.

#### Benzo-[$\alpha$]-pyrene Cytotoxic Doses Removed

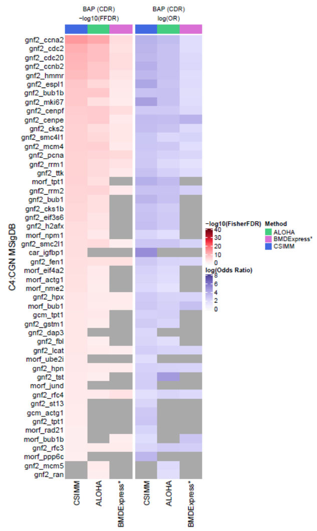


**Fig 38. BAP AD** Heatmaps of the significant Cancer Gene Neighborhood gene sets comparing the most statistically significant clusters, displaying negative log-scale Fisher FDR and log odds ratio, respectively. Gray indicates which gene sets were not enriched for a given clustering approach.

#### 4.7 Gene Set Level Benchmark Dose (BMD) Coherence

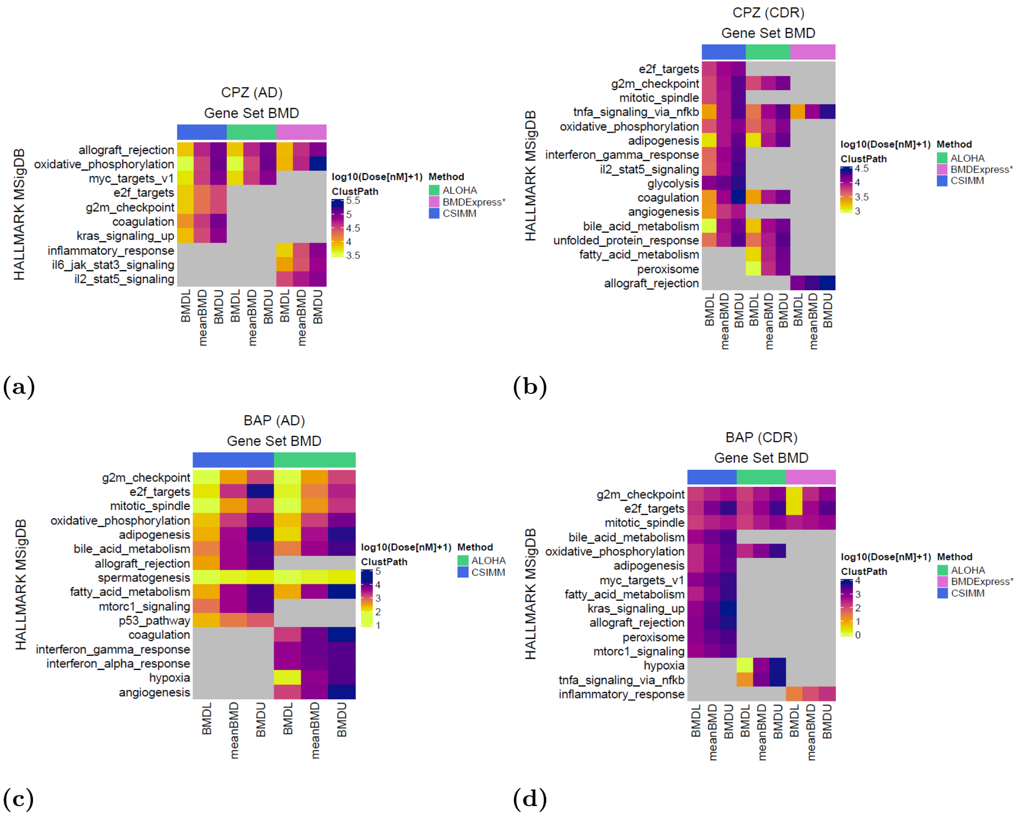


**Fig 39.** Heatmaps comparing the posterior mean BMD and 95% credible intervals (i.e. BMDL and BMDU) estimates (columns) for enriched Hallmark gene sets [10–12] (rows) across the three clustering methods. The BMDs are estimated using core genes in the most relevant cluster, as defined in §0.3.3, mapping to the respective gene sets. (a) Chlorpromazine with all experimental dose groups included. (b) Chlorpromazine with cytotoxic dose groups removed. (c) Benzo-[$\alpha$]-pyrene with all experimental dose groups. (d) Benzo-[$\alpha$]-pyrene with cytotoxic dose groups removed. Gray blocks within the heatmap indicate which gene sets are not enriched for a clustering approach. Bars on above the columns indicate the various clustering approaches, i.e. CSIMM (blue), ALOHA (green), and BMDExpress clustering (magenta), described in §0.2.

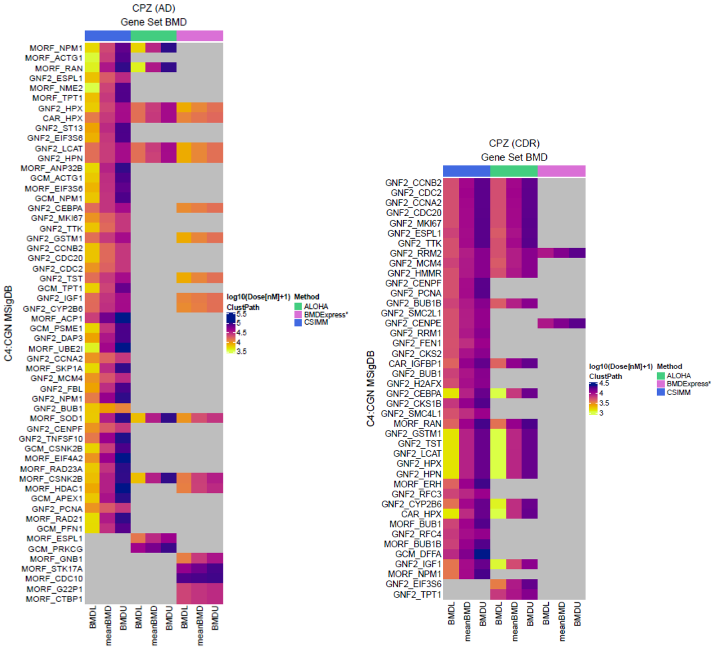


**(a)** **(b)**

**Fig 40.** Heatmaps comparing the posterior mean BMD and 95% credible interval (i.e. BMDL and BMDU) estimates (columns) for enriched Cancer Gene Neighborhoods (CGN) gene sets [10–12] (rows) across the three clustering methods. The BMDs are estimated using core genes in the most relevant cluster, as defined in §0.3.3, mapping to the respective gene sets. (a) Chlorpromazine with all experimental dose groups included. (b) Chlorpromazine with cytotoxic dose groups removed. Gray blocks within the heatmap indicate which gene sets are not enriched for a clustering approach. Bars on above the columns indicate the various clustering approaches, i.e. CSIMM (blue), ALOHA (green), and BMDExpress clustering (magenta), described in §0.2.

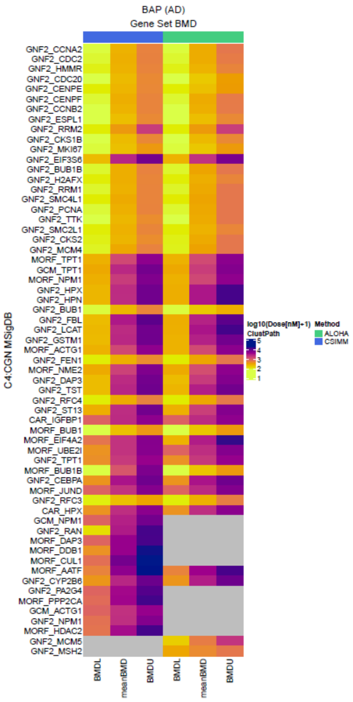


**(a)**

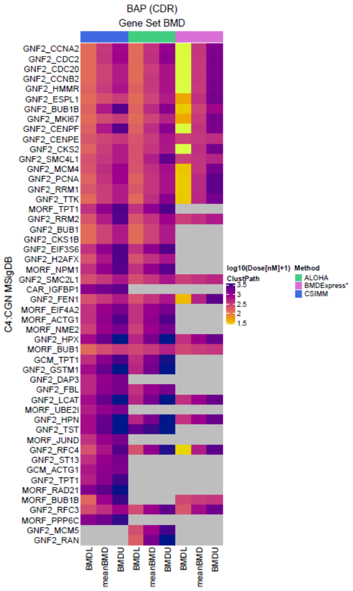


**(b)**

**Fig 41.** Heatmaps comparing the posterior mean BMD and 95% credible intervals (i.e. BMDL and BMDU) estimates (columns) for enriched Cancer Gene Neighborhoods (CGN) gene sets [10–12] (rows) across the three clustering methods. The BMDs are estimated using core genes in the most relevant cluster, as defined in §0.3.3, mapping to the respective gene sets. (a) Benzo-[$\alpha$]-pyrene with all experimental dose groups included. (b) Benzo-[$\alpha$]-pyrene with cytotoxic dose groups removed. Gray blocks within the heatmap indicate which gene sets are not enriched for a clustering approach. Bars on above the columns indicate the various clustering approaches, i.e. CSIMM (blue), ALOHA (green), and BMDExpress clustering (magenta), described in §0.2.

#### 4.8 Core Gene Comparison Tables

The following tables contain information about the core probes used to estimate the gene-set level BMDs in Fig 7.

**Chlorpromazine All Doses**

| BMDE genes | CSIMM genes | ALOHA genes | BMDExpress genes |
|---|---|---|---|
| ATF5_501, AURKA_586, CCND1_1062, CCNF_1076, CDC7_19277, CENPF_1252, CKS1B_1362, CKS2_19000, CTCF_19656, DBF4_20417, E2F2_2009, G3BP1_2542, H2AFX_26072, H2AFZ_2864, HSPA8_3142, MAD2L1_3930, MAPK14_4000, MARCKS_4021, MT2A_4334, MTF2_4342, ORC6_4832, PBK_19921, PLK1_5203, PRMT5_28286, PTTG1_5623, RACGAP1_19398, RAD21_5698, SMAD3_27880, SRSF10_24587, STIL_15150, STMN1_6857, TFDP1_7045, TGFB1_27949, TOP2A_7277, TPX2_7308 | CDC20_1157, HMMR_3040, KIF2C_3603, MAD2L1_3930, TOP2A_7277, TPX2_7308 | | |

**Table 6.** Core probes from the most significant cluster(s) enriched for the Hallmark gene set *"G2M Checkpoint"* in each clustering approach and BMDExpress use to estimate the gene set level BMD estimate and interval for the **CPZ-AD** dataset.

| BMDE genes | CSIMM genes | ALOHA genes | BMDExpress genes |
|---|---|---|---|
| ANP32E_26838, ATAD2_22038, AURKA_586, BRCA1_757, BRMS1L_22068, BUB1B_800, CCP110_26981, CDCA3_19307, CDCA8_1185, CDKN2A_28291, CKS1B_1362, CKS2_19000, CTCF_19656, CTPS1_16302, DCK_1776, DEK_1841, DE-PDC1_16570, DEPDC1_27104, DLGAP5_1882, H2AFX_26072, H2AFZ_2864, KIF18B_3598, LYAR_12352, MAD2L1_3930, MSH2_19521, NME1_27589, NOP56_4635, ORC6_4832, PAICS_4908, PCNA_4987, PLK1_5203, PSIP1_27736, PSIP1_5500, PTTG1_5623, RACGAP1_19398, RAD21_5698, RAD50_12781, RAD51C_5708, RAN_5725, RFC2_5805, RPA3_5906, SLBP_20689, SPAG5_17378, SPC25_6704, STMN1_6857, TIPIN_11359, TOP2A_7277, UBE2T_7492 | CDC20_1157, HMMR_3040, KIF2C_3603, MAD2L1_3930, SPC25_6704, TOP2A_7277 | | |

**Table 7.** Core probes from the most significant cluster(s) enriched for the Hallmark gene set *"E2F Targets"* in each clustering approach and BMDExpress use to estimate the gene set level BMD estimate and interval for the **CPZ-AD** dataset.

| BMDE genes | CSIMM genes | ALOHA genes | BMDExpress genes |
|---|---|---|---|
| ACADVL_20681, | AIFM1_183, | ACAA1_48, | ACADVL_20681, |
| ACAT1_22692, | ATP5B_26124, | AIFM1_183, | ACAT1_22692, |
| AIFM1_183, | ATP5C1_17193, | ATP5B_26124, | AIFM1_183, |
| ATP1B1_527, | ATP6AP1_14351, | ATP5C1_17193, | ATP1B1_527, |
| ATP5B_26124, | ATP6V0B_562, | ATP6AP1_14351, | ATP5B_26124, |
| ATP5C1_17193, | ATP6V0E1_23077, | ATP6V0B_562, | ATP5C1_17193, |
| ATP6AP1_14351, | ATP6V1D_574, | ATP6V0E1_23077, | ATP6AP1_14351, |
| ATP6V0E1_23077, | COX8A_19058, | ATP6V1D_574, | ATP6V0E1_23077, |
| ATP6V1D_574, | CYCS_1692, | COX8A_19058, | ATP6V1D_574, |
| BAX_28222, | DECR1_1831, | CYC1_27087, | BAX_28222, |
| CASP7_985, | ECH1_2022, | CYCS_1692, | CASP7_985, |
| CYCS_1692, | ECI1_22689, | DECR1_1831, | CYCS_1692, |
| DLAT_1875, | ETFB_2245, | ECH1_2022, | DLAT_1875, |
| ECH1_2022, | GPX4_17664, | ECI1_22689, | ECH1_2022, |
| ECI1_22689, | LDHA_3766, | ETFB_2245, | ECI1_22689, |
| ETFA_2244, | MDH1_4087, | GLUD1_2678, | ETFA_2244, |
| ETFB_2245, | MRPL15_4264, | GPX4_17664, | ETFB_2245, |
| GLUD1_2678, | NDUFA1_4514, | LDHA_3766, | GLUD1_2678, |
| GPI_2753, | NDUFB3_18124, | MDH1_4087, | GPI_2753, |
| GPX4_17664, | NDUFV2_4530, | MRPL15_4264, | GPX4_17664, |
| HSD17B10_3112, | PDHX_5033, | NDUFA1_4514, | HSD17B10_3112, |
| IDH3A_15224, | PRDX3_27724, | NDUFB3_18124, | IDH3A_15224, |
| MDH1_4087, | SDHB_6173, | NDUFV2_4530, | MDH1_4087, |
| MDH2_20964, | SLC25A4_6406, | PDHX_5033, | MDH2_20964, |
| MRPL15_4264, | SLC25A5_20900, | PRDX3_27724, | MRPL15_4264, |
| NDUFA1_4514, | SLC25A6_14758, | SDHB_6173, | NDUFA1_4514, |
| NDUFB3_18124, | SUPV3L1_6897, | SLC25A4_6406, | NDUFB3_18124, |
| NDUFV2_4530, | UQCRC1_13094, | SLC25A5_20900, | NDUFV2_4530, |
| PDHB_13121, | UQCRFS1_7566, | SLC25A6_14758, | PDHB_13121, |
| PDHX_5033, | VDAC1_7642, | SUPV3L1_6897, | PDHX_5033, |
| POR_23019, | VDAC2_28051 | UQCRC1_13094, | POR_23019, |
| PRDX3_27724, | | UQCRFS1_7566, | PRDX3_27724, |
| SDHB_6173, | | VDAC1_7642, | SDHB_6173, |
| SLC25A4_6406, | | VDAC2_28051 | SLC25A4_6406, |
| SLC25A5_20900, | | | SLC25A5_20900, |
| SLC25A6_14758, | | | SLC25A6_14758, |
| SUCLA2_6870, | | | SUCLA2_6870, |
| UQCRC1_13094, | | | UQCRC1_13094, |
| UQCRQ_7569, | | | UQCRQ_7569, |
| VDAC1_7642, | | | VDAC1_7642, |
| VDAC2_11431, | | | VDAC2_11431, |
| VDAC2_28051, | | | VDAC2_28051, |
| VDAC2_28052 | | | VDAC2_28052 |

**Table 8.** Core probes from the most significant cluster(s) enriched for the Hallmark gene set *"Oxidative Phosphorylation"* in each clustering approach and BMDExpress use to estimate the gene set level BMD estimate and interval.

| BMDE genes | CSIMM genes | ALOHA genes | BMDExpress genes |
|---|---|---|---|
| ACE_26794,<br>ADAM8_12335,<br>CA2_908,<br>CFH_27007,<br>CTSS_25157,<br>DOCK2_20159,<br>DUSP6_1993,<br>ELTD1_21057,<br>ELTD1_28345,<br>EMP1_18941,<br>ENG_13441,<br>ETS1_27185,<br>ETV1_2251,<br>ETV1_27186,<br>FCER1G_11058,<br>G0S2_13343,<br>GLRX_2674,<br>IGFBP3_3270,<br>IKZF1_3293,<br>IL1B_3325,<br>IL2RG_15624,<br>INHBA_21251,<br>IRF8_20223,<br>KIF5C_3612,<br>LAPTM5_23860,<br>SERPINA3_17487,<br>SNAP25_6607,<br>TLR8_16795 | ACE_26793,<br>DUSP6_1993,<br>ELTD1_21057,<br>ELTD1_28345,<br>ENG_13441,<br>ETV1_2251,<br>IL2RG_15624,<br>INHBA_21251,<br>KIF5C_3612,<br>LY96_18964,<br>MAP7_3988,<br>TLR8_16795 | | |

**Table 9.** Core probes from the most significant cluster(s) enriched for the Hallmark gene set *"KRAS Signaling Up"* in each clustering approach and BMDExpress use to estimate the gene set level BMD estimate and interval.

**Benzo-[$\alpha$]-pyrene All Doses**

| BMDE genes | CSIMM genes | ALOHA genes | BMDExpress genes |
|---|---|---|---|
| AURKA_586, AURKB_587, BIRC5_709, CCNF_1076, CDC25B_1161, CENPA_1247, CENPF_1252, CHEK1_1288, CKS1B_1362, CKS2_19000, EGF_2055, EXO1_17006, EZH2_2282, FBXO5_10555, G3BP1_2542, H2AFX_26072, HIF1A_2945, HMMR_3040, KIF15_3595, KIF23_14714, KIF2C_3603, KIF4A_28251, MAD2L1_3930, MCM3_4072, MKI67_28355, MT2A_4334, MTF2_4342, NOTCH2_18421, ORC6_4832, PLK1_5203, PRC1_14588, RACGAP1_19398, RAD21_5698, RAD54L_14089, RPA2_5905, SLC7A5_13711, SMAD3_27880, SMARCC1_6575, SMC1A_6580, SMC2_12897, STIL_15150, TACC3_15729, TTK_7394, UBE2C_7468 | AURKB_587, CCNA2_1050, CCNF_1076, CDC20_1157, CDC45_26987, CDC6_1177, CDK1_1196, CENPF_1252, CKS1B_1362, EZH2_2282, FBXO5_10555, HMMR_3040, KIF11_3588, KIF23_14714, KIF2C_3603, MKI67_28355, NEK2_20563, NUSAP1_4789, PLK1_5203, PRC1_14588, SMC1A_6580, SMC4_6586, TACC3_15729, TPX2_7308 | AURKB_587, CCNA2_1050, CCNF_1076, CDC20_1157, CDC45_26987, CDC6_1177, CDK1_1196, CENPF_1252, CKS1B_1362, CKS2_19000, EZH2_2282, FBXO5_10555, HMMR_3040, KIF11_3588, KIF23_14714, KIF2C_3603, MCM3_4072, MKI67_28355, MT2A_4334, NEK2_20563, NUSAP1_4789, ORC6_4832, PLK1_5203, PRC1_14588, RACGAP1_19398, RAD54L_14089, SMC1A_6580, SMC4_6586, TACC3_15729, TPX2_7308 | |

**Table 10.** Core probes from the most significant cluster(s) enriched for the Hallmark gene set *"G2M Checkpoint"* in each clustering approach and BMDExpress use to estimate the gene set level BMD estimate and interval for the **BAP-AD** dataset.

| BMDE genes | CSIMM genes | ALOHA genes | BMDExpress genes |
|---|---|---|---|
| ANLN_12002, ARHGEF2_423, AURKA_586, BCL2L11_26886, BIRC5_709, CDC42_1168, CENPF_1252, CEP57_1257, DLGAP5_1882, FBXO5_10555, HDAC6_2901, KIF15_3595, KIF23_14714, KIF2C_3603, KIF4A_28251, NCK1_4491, NOTCH2_18421, OPHN1_28310, PCM1_4980, PLK1_5203, PRC1_14588, PXN_5642, RACGAP1_19398, SMC1A_6580, SMC3_6583, SOS1_10682, SPTAN1_6727, TTK_7394 | ANLN_12002, CDK1_1196, CENPF_1252, FBXO5_10555, KIF11_3588, KIF23_14714, KIF2C_3603, NEK2_20563, NUSAP1_4789, PLK1_5203, PRC1_14588, SMC1A_6580, SMC4_6586, TPX2_7308 | ANLN_12002, CDK1_1196, CENPF_1252, FBXO5_10555, KIF11_3588, KIF23_14714, KIF2C_3603, NEK2_20563, NUSAP1_4789, PLK1_5203, PRC1_14588, RACGAP1_19398, SMC1A_6580, SMC4_6586, TPX2_7308 | |

**Table 11.** Core probes from the most significant cluster(s) enriched for the Hallmark gene set *"Mitotic Spindle"* in each clustering approach and BMDExpress use to estimate the gene set level BMD estimate and interval for the **BAP-AD** dataset.

| BMDE genes | CSIMM genes | ALOHA genes | BMDExpress genes |
|---|---|---|---|
| ACAA1_48, ACADVL_20681, ALAS1_28273, ATP5J_13202, ATP6AP1_14351, ATP6V1D_574, DLAT_1875, ECI1_22689, ETFA_2244, MDH1_4087, NDUFA1_4514, NDUFB3_18124, NDUFV2_4530, POR_23019, PRDX3_27724, SDHB_6173, SUPV3L1_6897, TIMM9_7106, VDAC1_7642, VDAC2_11431 | ATP1B1_527, ATP5C1_17193, COX6B1_1519, CYC1_27087, CYCS_1692, DLAT_1875, GPI_2753, GPX4_17664, LDHA_3766, MAOB_13383, MDH2_20964, MRPL15_4264, NDUFA1_4514, NDUFA4_14125, NDUFB3_18124, NDUFV2_4530, NNT_4618, SDHB_6173, SLC25A4_6406, SLC25A6_14758, SUCLA2_6870, UQCRQ_7569 | ACAA1_48, ATP1B1_527, COX6B1_1519, CYC1_27087, CYCS_1692, DECR1_1831, DLAT_1875, GPI_2753, GPX4_17664, LDHA_3766, MDH2_20964, NDUFA1_4514, NDUFA4_14125, NDUFB3_18124, NDUFV2_4530, NNT_4618, SDHB_6173, SLC25A4_6406, SLC25A5_20900, SLC25A6_14758, SUCLA2_6870, UQCRQ_7569 | |

**Table 12.** Core probes from the most significant cluster(s) enriched for the Hallmark gene set *"Oxidative Phosphorylation"* in each clustering approach and BMDExpress use to estimate the gene set level BMD estimate and interval for the **BAP-AD** dataset.

| BMDE genes | CSIMM genes | ALOHA genes | BMDExpress genes |
|---|---|---|---|
| APP_367, ATF3_499, BLCAP_713, BTG2_13191, CASP1_26966, CCP110_26981, DDIT3_16736, FAS_2345, FOXO3_2481, GPX2_11201, IKBKAP_3288, IL1A_22714, MDM2_23384, MDM2_27496, PHLDA3_18856, RAD51C_5708, RALGDS_5721, RNF19B_24383, SAT1_17598, SFN_6269, SLC7A11_14100, TM7SF3_22780, TP53_7287, TRIAP1_21159 | AEN_18198, CASP1_26966, CCP110_26981, DDB2_1798, FAS_2345, GADD45A_2569, PPM1D_25317, PRKAB1_14557, PROCR_24511, SDC1_6160, XPC_28078 | | |

**Table 13.** Core probes from the most significant cluster(s) enriched for the Hallmark gene set *"P53 Pathway"* in each clustering approach and BMDExpress use to estimate the gene set level BMD estimate and interval for the **BAP-AD** dataset.
